# Distributed representations of behavior-derived object dimensions in the human visual system

**DOI:** 10.1101/2023.08.23.553812

**Authors:** O. Contier, C.I. Baker, M.N. Hebart

## Abstract

Object vision is commonly thought to involve a hierarchy of brain regions processing increasingly complex image features, with high-level visual cortex supporting object recognition and categorization. However, object vision supports diverse behavioral goals, suggesting basic limitations of this category-centric framework. To address these limitations, we mapped a series of dimensions derived from a large-scale analysis of human similarity judgments directly onto the brain. Our results reveal broadly distributed representations of behaviorally-relevant information, demonstrating selectivity to a wide variety of novel dimensions while capturing known selectivities for visual features and categories. Behavior-derived dimensions were superior to categories at predicting brain responses, yielding mixed selectivity in much of visual cortex and sparse selectivity in category-selective clusters. This framework reconciles seemingly disparate findings regarding regional specialization, explaining category selectivity as a special case of sparse response profiles among representational dimensions, suggesting a more expansive view on visual processing in the human brain.

## Introduction

A central goal of visual neuroscience is to understand how the brain encodes and represents rich information about objects, allowing us to make sense of our visual world and act on it in meaningful ways. A widely studied and influential account posits that one central function of the visual system is to recognize objects by organizing them into distinct categories (DiCarlo et al., 2012; Goodale & Milner, 1992; Marr, 2010; Mishkin & Ungerleider, 1982). According to this view, early visual cortex serves to analyze incoming visual information by representing basic visual features (Hubel & Wiesel, 1962), which are then combined into more and more complex feature combinations, until higher-level visual regions in the occipitotemporal cortex and beyond support the recognition of object identity and category (DiCarlo et al., 2012). In line with this view, a number of category-selective clusters have been identified in occipitotemporal cortex that respond selectively to specific object classes such as faces, scenes, body parts, tools, or text (Downing & Kanwisher, 2010; Epstein & Kanwisher, 1998; Kanwisher et al., 1997; Kanwisher & Yovel, 2006; A. Martin et al., 1996; Puce et al., 1996). The functional significance of these regions is underscored by studies demonstrating that object category and identity as well as performance in some behavioral tasks can be read out from activity in occipitotemporal cortex (Carlson et al., 2014; Cohen et al., 2017; Hung et al., 2005; Ritchie et al., 2015; Ritchie & Carlson, 2016; Singer et al., 2023) and that lesions to these regions can lead to selective deficits in object recognition abilities (Kanwisher & Barton, 2011; Konen et al., 2011; Moro et al., 2008; Schiltz et al., 2006; Wada & Yamamoto, 2001).

Despite the importance of object categorization and identification as crucial goals of object vision, it has been argued that these functions alone are insufficient for capturing how our visual system allows us to make sense of the objects around us (Bracci & Op de Beeck, 2023). A more comprehensive understanding of object vision should account for the rich meaning and behavioral relevance associated with individual objects beyond discrete labels. This requires incorporating the many visual and semantic properties of objects that underlie our ability to make sense of our visual environment, perform adaptive behaviors, and communicate about our visual world (Bracci & Op de Beeck, 2023; Cox, 2014; Krakauer et al., 2017; Kravitz et al., 2013; Peelen & Downing, 2017). Indeed, others have proposed that visual cortex is organized based on continuous dimensions reflecting more general object properties, such as animacy (Bao et al., 2020; Caramazza & Shelton, 1998; Konkle & Caramazza, 2013; Kriegeskorte, 2009), real-world size (Konkle & Caramazza, 2013; Konkle & Oliva, 2012), aspect ratio (Bao et al., 2020; Coggan & Tong, 2023), or semantics (Huth et al., 2012). These and other continuous dimensions reflect behaviorally-relevant information that offers a more fine-grained account of object representations than discrete categorization and recognition alone. This dimensional view suggests a framework in which visual cortex is organized based on topographic tuning to specific dimensions that extends beyond category-selective clusters. Under this framework, category-selective clusters may emerge from a more general organizing principle (Haxby et al., 2001; Huth et al., 2012; Mahon & Caramazza, 2011; A. Martin, 2007; Op de Beeck et al., 2008), reflecting cortical locations where these tuning maps encode feature combinations tied to specific object categories (Arcaro & Livingstone, 2021; Huth et al., 2012; Op de Beeck et al., 2008). Yet, while previously proposed dimensions have been shown to partially reflect activity patterns in category-selective clusters (Andrews et al., 2010; Coggan et al., 2016, 2019; Nasr et al., 2014; Nasr & Tootell, 2012; Rice et al., 2014), they cannot account fully for the response profile and are largely inferior to category-selectivity in explaining the functional selectivity of human visual cortex for objects (Downing et al., 2006; Yargholi & Op de Beeck, 2023).

To move beyond the characterization of individual behavioral goals underlying both the discrete category-centric and the continuous dimensional view and to comprehensively map a broad spectrum of behaviorally-relevant representations, one powerful approach is to link object responses in visual cortex to judgments about the perceived similarity between objects (Charest et al., 2014; Cichy et al., 2019; Magri & Konkle, 2019; Mur et al., 2013). Indeed, perceived similarity serves as a common proxy of mental object representations underlying various behavioral goals, as the similarity relation between objects conveys much of the object knowledge and behavioral relevance across diverse perceptual and conceptual criteria (Ashby & Perrin, 1988; Edelman, 1998; Hebart et al., 2020; Nosofsky, 1984; Shepard, 1987). As such, perceived similarity is ideally suited for revealing behaviorally-relevant representational dimensions and how these dimensions are reflected in cortical patterns of brain activity.

To uncover the nature of behaviorally-relevant selectivity underlying similarity judgments in human visual cortex, in the present study we paired functional MRI responses to thousands of object images (Hebart et al., 2023) with core representational dimensions derived from a dataset of millions of human similarity judgments. In contrast to much previous research that has focused on a small number of hypothesis-driven dimensions or that used small, selective image sets (Almeida et al., 2023; Bracci & Op de Beeck, 2016; Charest et al., 2014; Cichy et al., 2019; Konkle & Caramazza, 2013; Kriegeskorte et al., 2008; Magri & Konkle, 2019; Mur et al., 2013), we carried out a comprehensive characterization of cortical selectivity in response to 66 representational dimensions identified in a data-driven fashion for 1,854 objects (Hebart et al., 2020; Zheng et al., 2019).

Moving beyond the view that mental object representations derived from similarity judgments are primarily mirrored in high-level visual cortex (Charest et al., 2014; Cichy et al., 2019; Hebart et al., 2023; Mur et al., 2013), we demonstrate that representations underlying core object dimensions are reflected throughout the entire visual cortex. Our results reveal that cortical tuning to these dimensions captures the functional topography of visual cortex and mirrors stimulus selectivity throughout the visual hierarchy. In this multidimensional representation, category selectivity stands out as a special case of sparse selectivity to a set of core representational object dimensions, while other parts of visual cortex reflect a more mixed selectivity. A direct model comparison revealed that continuous object dimensions provide a better model of brain responses than categories across the visual system, suggesting that dimension-related tuning maps offer more explanatory power than a category-centric framework. Together, our findings reveal the importance of behavior-derived object dimensions for understanding the functional organization of the visual system and offer a broader, comprehensive view of object representations that bridges the gap between regional specialization and domain-general topography.

## Results

We first aimed at mapping core representational object dimensions to patterns of brain activity associated with visually-perceived objects. To model the neural representation of objects while accounting for their large visual and semantic variability (Groen et al., 2017; Naselaris et al., 2021), we used the THINGS-data collection (Hebart et al., 2023), which includes densely sampled fMRI data for thousands of naturalistic object images from 720 semantically diverse objects, as well as 4.7 million behavioral similarity judgments of these objects (Fig. 1).

**Fig. 1.**
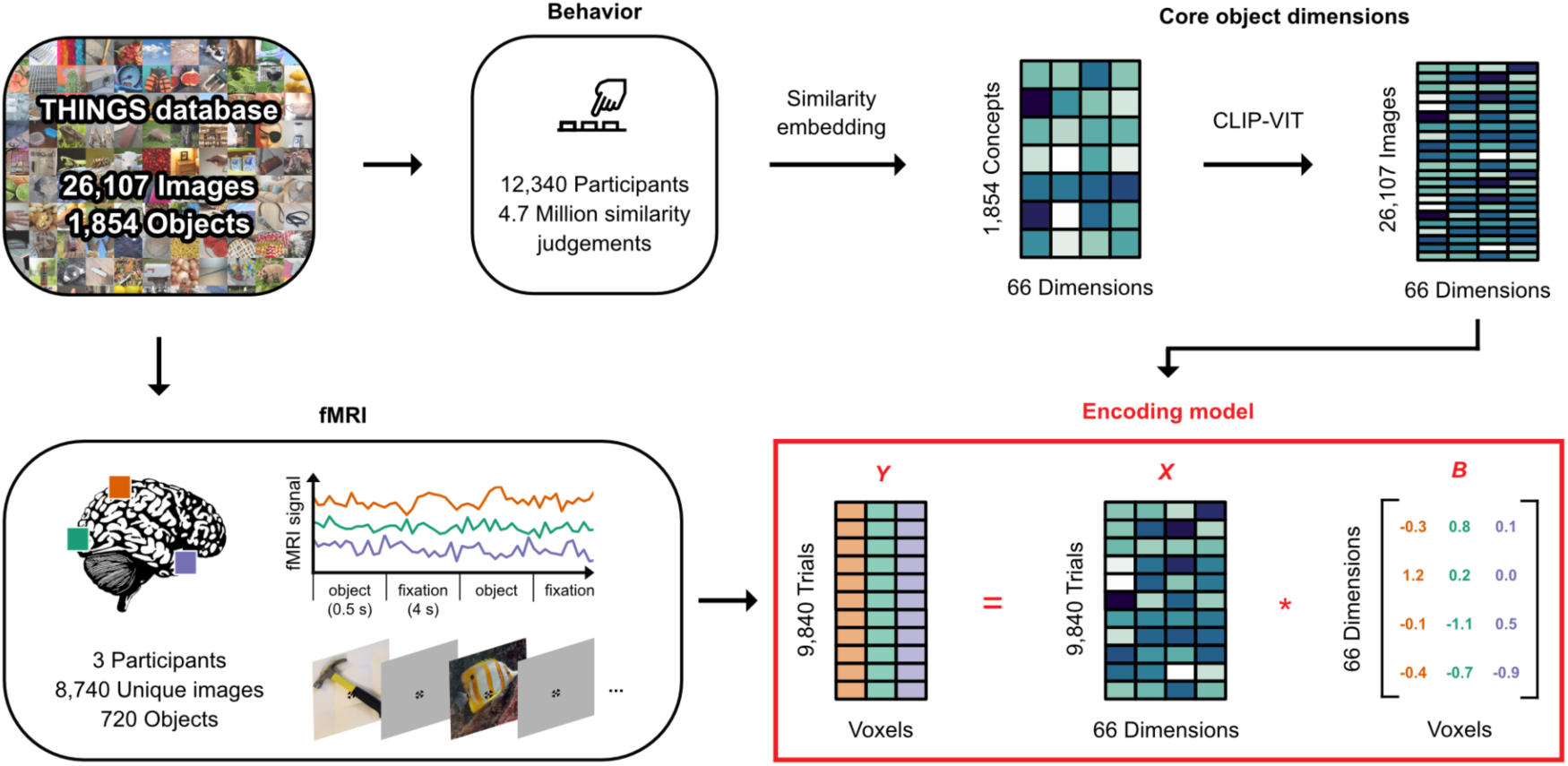
Overview: An fMRI encoding model of object dimensions underlying human similarity judgements. We linked core representational dimensions capturing the behavioral relevance of objects to spatially resolved neural responses to thousands of object images. For this, we used the THINGS-data collection (Hebart et al., 2023) which includes fMRI and behavioral responses to objects from the THINGS object concept and image database (Hebart et al., 2019). The behavioral data was used to train a computational model of core object dimensions underlying human similarity judgements to different object concepts. We extended this embedding to the level of individual object images based on the computer vision model CLIP-VIT (Radford et al., 2021). The fMRI data comprises three participants who each saw 8,740 unique object images. We used an encoding model of the object dimension embedding to predict fMRI responses to each image in each voxel. The estimated encoding model weights reflect the tuning of each voxel to each object dimension. *X*, *B*, and *Y* denote the design matrix, regression weights, and outcome of the encoding mode, respectively.

As core object dimensions, we used a recent similarity embedding of behavior-derived object dimensions, which underlie the perceived similarity of 1,854 object concepts (Hebart et al., 2020, 2023). In this embedding, each object image is characterized by 66 dimensions derived from the human similarity judgments in an odd-one-out task. We chose this embedding for several reasons: First, it provides highly reproducible dimensions that together are sufficient for capturing single trial object similarity judgments close to the noise ceiling. Second, the use of an odd-one-out task supports the identification of the minimal information required to distinguish between different objects, and as such is sensitive not only to conceptual information, such as high-level category (e.g., "is an animal"), but also to key visual-perceptual distinctions (e.g., "is round"). Thus, the object dimensions capture behaviorally-relevant information, in that they support the key factors underlying arbitrary categorization behavior and as such underlie our ability to make sense of our visual world, to generalize, structure our environment, and to communicate our knowledge. Indeed, the object dimensions capture external behavior such as high-level categorization and typicality judgements, underscoring their potential explanatory value as a model of neural responses to objects (Hebart et al., 2020). Third, the object dimensions are easily interpretable, thus simplifying the interpretation of neural activity patterns in relation to individual dimensions.

The fMRI dataset covers 8,740 unique images from 720 categories presented to three participants (2 female) over the course of 12 sessions (Hebart et al., 2023). Given that the behavioral similarity embedding was trained only on one image per each of the 1,854 THINGS categories, these dimensions may only partially capture the visual richness of the entire image set, which may affect the potential for predicting image-wise brain responses. To address this challenge, we fine-tuned the artificial neural network model CLIP-VIT (Radford et al., 2021) to directly predict object dimensions for the 8,740 images in our fMRI dataset. This model has previously been shown to provide a good correspondence to behavioral (Muttenthaler et al., 2023; Muttenthaler & Hebart, 2021) and brain data (Conwell et al., 2023; Wang et al., 2023), indicating its potential for providing accurate image-wise estimates of behavior-derived object dimensions. Indeed, this prediction approach led to highly accurate cross-validated predictions of object similarity (Kaniuth et al., 2024) and consistent improvements in BOLD signal predictions for all 66 dimensions (Suppl. Fig. 1).

### Core object dimensions are reflected in widespread fMRI activity patterns throughout the human visual system

To test how these dimensions were expressed in voxel-wise brain responses, we fit an fMRI encoding model which predicts spatially resolved brain responses based on a weighted sum of these object dimensions. This allowed us to map out the contribution of the dimensions to the measured signal and thus link interpretable behavior-derived dimensions to patterns of brain activity.

Across all 66 object dimensions, our results revealed a widely distributed cortical representation of these dimensions that spans much of visual cortex and beyond (Fig. 2). The spatial extent of these effects was highly similar across all three subjects, underscoring the generality of these findings. We also tested the replicability of these results on an independent fMRI dataset (Chang et al., 2019), revealing a similarly extensive representation of the object dimensions (Suppl. Fig. 2). Please note that, in the following, we use the terms "widespread" and "distributed" interchangeably and do not refer to a distributed representational coding scheme or the presence of continuous gradients but rather to responses that are not locally confined.

**Fig. 2.**
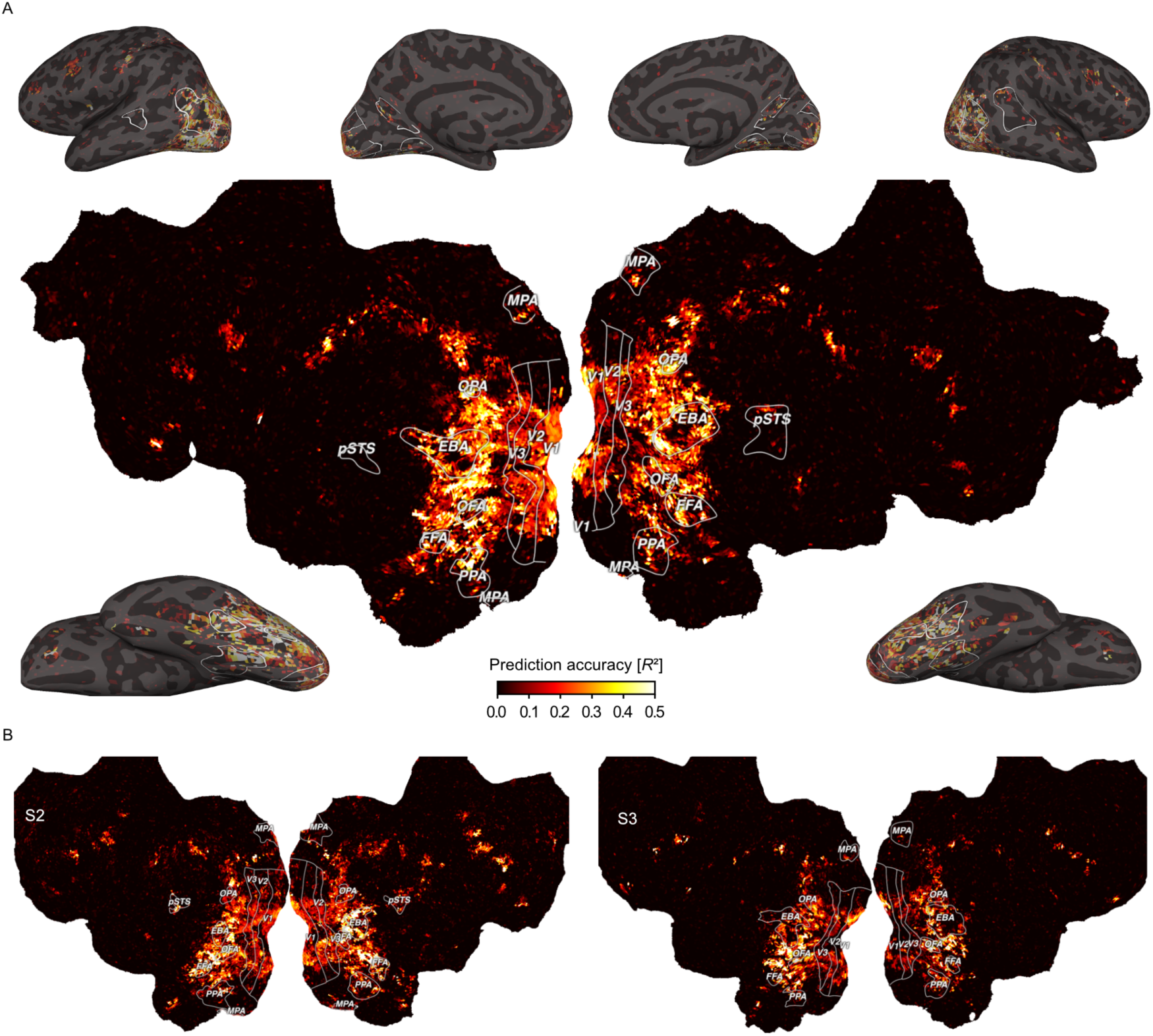
Prediction accuracy of the fMRI voxel-wise encoding model based on 66 core object dimensions. Colors indicate the proportion of explained variance (noise ceiling corrected R^2^) of held-out data in a 12-fold between-session cross-validation. White outlines indicate regions of interests defined in separate localizer experiments: FFA: Fusiform face area; OFA: Occipital face area; pSTS: Posterior superior temporal sulcus; EBA: Extrastriate body area; PPA: Parahippocampal place area; OPA: Occipitoparietal place area; MPA: Medial place area; V1-V3: Primary to tertiary visual cortex. A. Prediction accuracy for one example subject (S1) visualized on a cortical flat map (center) and inflated views of the cortical surface (corners). B. Results for the other two subjects visualized on cortical flat maps.

Prediction accuracies peaked not only in lateral occipital and posterior ventral temporal regions, but also reached significant values in early visual, dorsal visual, and frontal regions (Suppl. Fig. 3). In contrast to previous work based on representational similarity analysis that found information about perceived similarity to be confined primarily to higher-level visual cortex (Charest et al., 2014; Cichy et al., 2019; Hebart et al., 2023; Magri & Konkle, 2019), our dimension-based approach revealed that behaviorally-relevant information about objects is much more distributed throughout the visual processing hierarchy, including the earliest cortical processing stages.

### Behavior-derived object dimensions reflect the functional topography of the human visual system

Having identified where information about perceived similarity is encoded, we next explored the spatial layout of each individual dimension underlying this representation. By using a voxel-encoding model of interpretable object dimensions, it is possible to inspect the cortical distribution of the weights of each regressor separately and interpret them in a meaningful fashion. This has two benefits. First, it allows us to probe to what degree behavior-derived dimensions alone can capture the known topography of visual cortex. Second, it allows us to identify novel topographic patterns across visual cortex. This provides important insights into how the topography of visual cortex reflects object information relevant to behavior and how functionally specialized regions are situated in this cortical landscape.

Visualizing the voxel-wise regression weights for each object dimension on the cortical surface (Fig. 3) revealed a clear correspondence between numerous dimensions and characteristic, known topographic patterns of the visual system. For example, the "animal-related" dimension mirrors the well established spoke-like tuning gradient for animate versus inanimate objects (Konkle & Caramazza, 2013), while dimensions like "head-related" and "body-part related" differentiate the regional selectivity for faces and body parts in the fusiform face area (FFA), occipital face area (OFA), and extrastriate body area (EBA), respectively (Downing & Kanwisher, 2010; Gauthier et al., 2000; Kanwisher & Yovel, 2006). Likewise, the implicit inclusion of natural scenes as object backgrounds revealed scene content-related dimensions (e.g. "house-/ furnishing-related", "transportation-/ movement-related", and "outdoors"), which were found to be associated with scene-selective brain regions such as parahippocampal place area (PPA), medial place area (MPA), and occipital place area (OPA) (Epstein et al., 2005; Epstein & Kanwisher, 1998; Grill-Spector, 2003; Hasson et al., 2003; O’Craven & Kanwisher, 2000; Silson et al., 2016). Our approach also independently identified a "food-related" dimension in areas adjacent to the fusiform gyrus, in line with recently reported clusters responding selectively to food stimuli (Jain et al., 2023; Khosla et al., 2022; Pennock et al., 2023). A dimension related to tools ("tool-related/handheld/elongated") also matched expected activation patterns in middle temporal gyrus (He et al., 2020; A. Martin et al., 1996; A. Martin & Weisberg, 2003). Further, dimensions related to low- to mid-level visual features (e.g. "grid/grating-related", "repetitive/spiky") reflected responses primarily in early visual cortex.

**Fig. 3.**
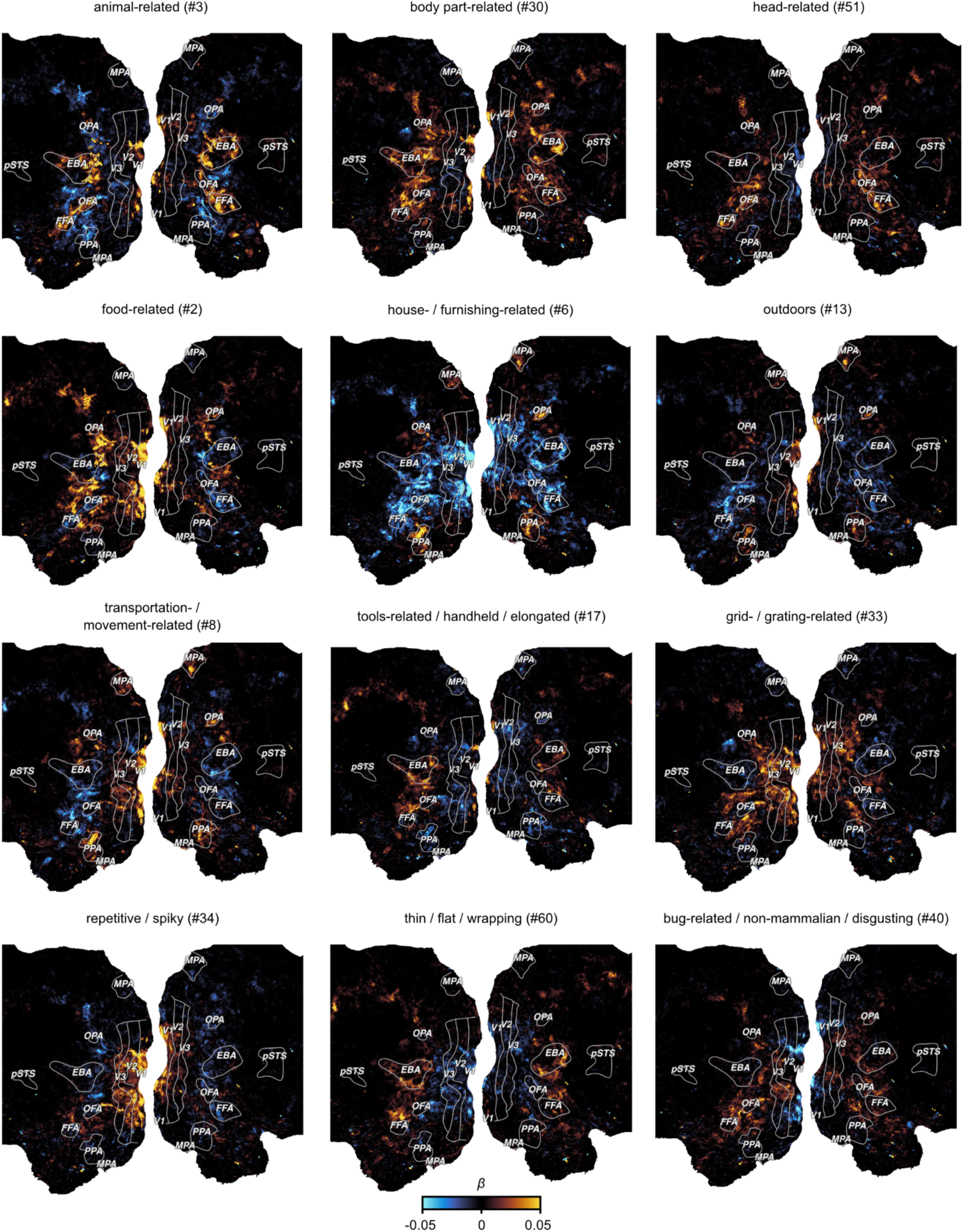
Functional tuning maps to individual object dimensions. The figure shows example maps for 12 out of the 66 dimensions for Subject S1. Each panel shows the encoding model weights for one object dimension projected onto the flattened cortical surface. Numbers in the subtitles show the dimension number in the embedding.

Beyond these established topographies, the results also revealed numerous additional topographic patterns. For example, one dimension reflected small, non-mammalian animals ("bug-related / non-mammalian / disgusting") that was clearly distinct from the "animal-related" dimension by lacking responses in face and body selective regions. Another dimension reflected a widely distributed pattern in response to thin, flat objects ("thin / flat / wrapping"). Thus, our approach allowed for the identification of candidate functional selectivities in visual cortex that might have gone undetected with more traditional approaches based on proposed categories or features (Downing et al., 2006; Khosla et al., 2022). Importantly, the functional topographies of most object dimensions were also found to be highly consistent across the three subjects in this dataset (Suppl. Fig. 4) and largely similar to participants of an independent, external dataset (Suppl. Fig. 2), suggesting that these topographies may reflect general organizing principles rather than idiosyncratic effects (Suppl. Fig. 4, Extended Data Fig. 1-6).

Together, our results uncover cortical maps of object dimensions underlying the perceived similarity between objects. These maps span extensive portions of the visual cortex, capturing topographic characteristics such as tuning gradients of object animacy, lower-level visual feature tuning in early visual cortex, and category-selective, higher-level regions while uncovering new candidate selectivities. Thus, these findings support an organizing principle where multiple, superimposing cortical tuning maps for core object properties collectively represent behaviorally-relevant information of objects.

### Cortical tuning to behavior-derived object dimensions explains regional functional selectivity

Having delineated the multidimensional topographic maps across visual cortex, we next honed in on individual brain regions to determine their functional selectivity as defined by their response tuning across these behavior-derived dimensions. To this end, we developed a high-throughput method to identify object images representative for specific brain regions. Specifically, we first determined a functional tuning profile across dimensions for each region of interest based on the region’s mean encoding model weights. Next, we identified images whose behavioral dimension profile best matched the functional tuning profile of the brain region. To this end, we used all 26,107 object images in the THINGS database (Hebart et al., 2019), most of which were unseen by participants, and assessed the cosine similarity between the dimension profiles of brain regions and images. This enabled us to rank over 26,000 images based on their similarity to a given brain region’s functional tuning profile.

Despite having been fitted solely on the 66-dimensional similarity embedding, our approach successfully identified diverse functional selectivities of visual brain regions (Fig. 4). For instance, the most representative images for early visual regions (V1, V2, V3) contained fine-scale, colorful, and repeating visual features, consistent with known representations of oriented edges and color in these areas (Hubel & Wiesel, 1968; Livingstone & Hubel, 1984). These patterns appeared more fine-grained in earlier (V1 or V2) compared to later retinotopic regions (hV4), potentially reflecting increased receptive field size along the retinotopic hierarchy (Kastner et al., 2001; Smith et al., 2001; Tootell et al., 1997). A similar finding is reflected in dimension selectivity profiles (Fig. 4), revealing increased color selectivity in hV4 compared to early retinotopic regions V1-V3 while yielding reductions in the "repetitive/spiky" dimension. Notably, tuning profiles in category-selective regions aligned with images of expected object categories: faces in face-selective regions (FFA, OFA), body parts in body-part selective regions (EBA), and scenes in scene-selective regions (PPA,

**Fig. 4.**
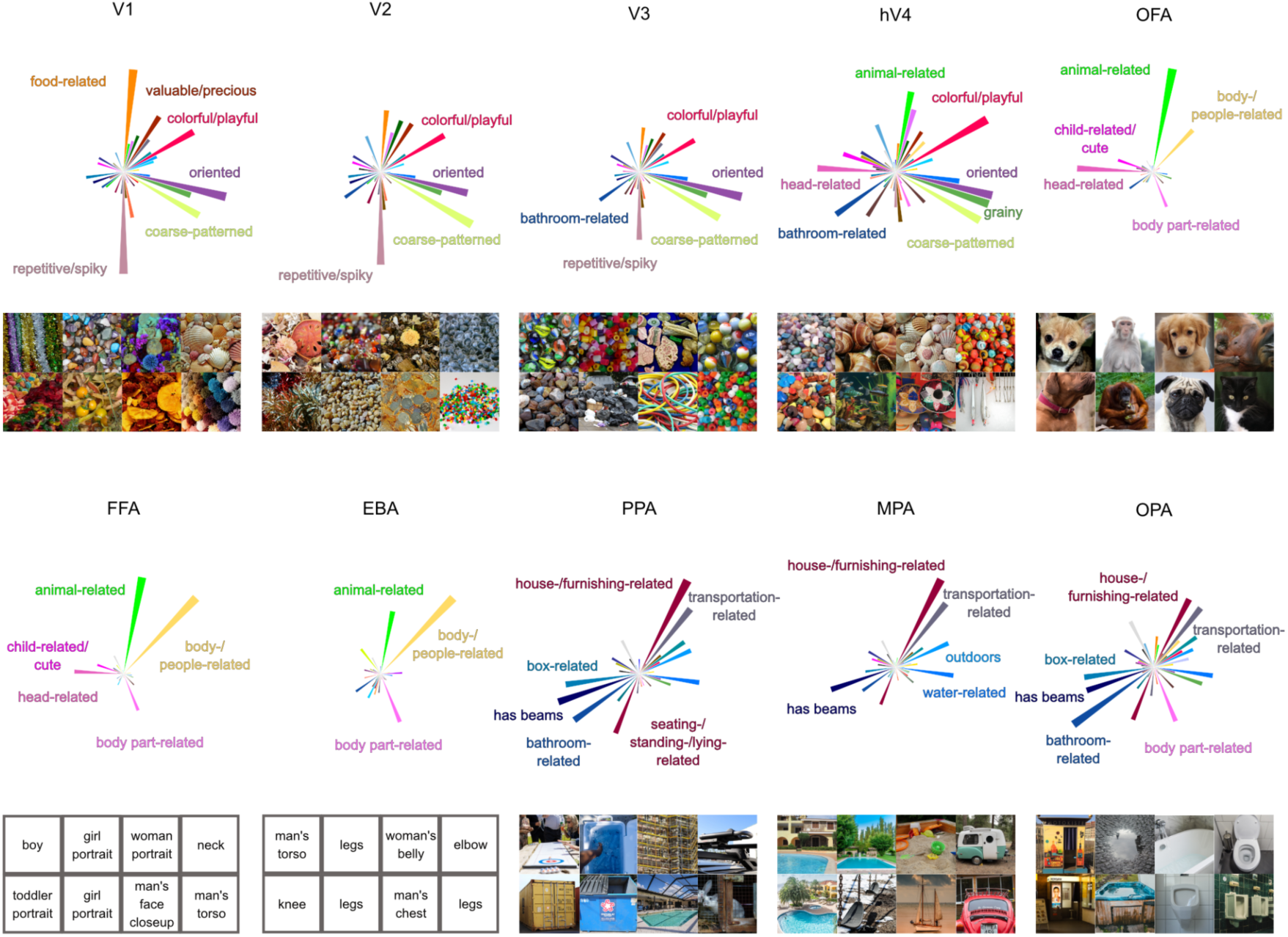
Regional tuning profiles across 66 object dimensions and representative images for selectivity of each region of interest in visual cortex. Rose plots indicate the magnitude of tuning for each object dimension in a given visual brain region. Image panels show 8 images with the most similar model representation to the regional tuning profile. For copyright reasons, all original images have been replaced with visually similar images. For identity protection reasons, images containing human faces and body parts have been replaced with verbal descriptions. Original images are available upon request.

OPA, MPA). Closer inspection of the tuning profiles revealed differences between regions that respond to the same basic object category, such as a stronger response to the "body-part related" dimension in OPA but not in other place-selective regions. Also, selectivity to faces (FFA, OFA) vs. body parts (EBA) appeared to be driven by the response magnitude to the "head-related" dimension, while tuning to the remaining dimensions was highly similar across these regions. Together, these findings demonstrate that the 66 object dimensions derived from behavior capture the selectivity across the visual processing hierarchy, highlighting the explanatory power of the dimensional framework for characterizing the functional architecture of the visual system.

### Category-selective brain regions are sparsely tuned to behavior-derived object dimensions

Given that dimensional tuning profiles effectively captured the selectivity of diverse visual regions, we asked what factors distinguish category-selective visual brain regions from non-category-selective regions in this dimensional framework. We reasoned that category-selectivity reflects a sparsely tuned representation, where activity in category-selective regions is driven by only a few dimensions, while non-category-selective regions reflect a more mixed selectivity, with activity related to a larger number of dimensions. In this way, functionally specialized, category-selective regions might stand-out as an extreme case of multidimensional tuning. As a consequence, this would also make it easier to identify category-selective regions due to their sparser selectivity.

To quantify this, we estimated a measure of sparseness over the encoding model weights in each voxel. Large sparseness indicates regions that are selective to very few dimensions, while lower sparseness indicates a dense representation in regions that respond broadly to diverse dimensions. Our results (Fig. 5A) indeed revealed sparser tuning in category-selective regions compared to other parts of the visual system. This effect was most pronounced in face and body part selective regions (FFA, OFA, EBA), with the sparsest tuning across all subjects. The face-selective posterior superior temporal sulcus exhibited particularly sparse representation in Subjects 1 and 2, while this region was not present in Subject 3 and, as expected, also yielded no increase in sparseness. Scene-selective regions (PPA, MPA, OPA) also exhibited sparseness, though the effects were more variable across subjects, which could arise from the representational dimensions being derived from objects within scenes, as opposed to isolated scene images without a focus on individual objects. Conversely, non-category-selective regions, such as early visual cortices, clearly exhibited dense representations. These findings suggest that category-selective regions, while responsive to multiple dimensions, may primarily respond to a small subset of behaviorally-relevant dimensions. Thus, in a multidimensional representational framework, category-selectivity may reflect a special case of sparse tuning within a broader set of distributed dimension tuning maps.

**Fig. 5.**
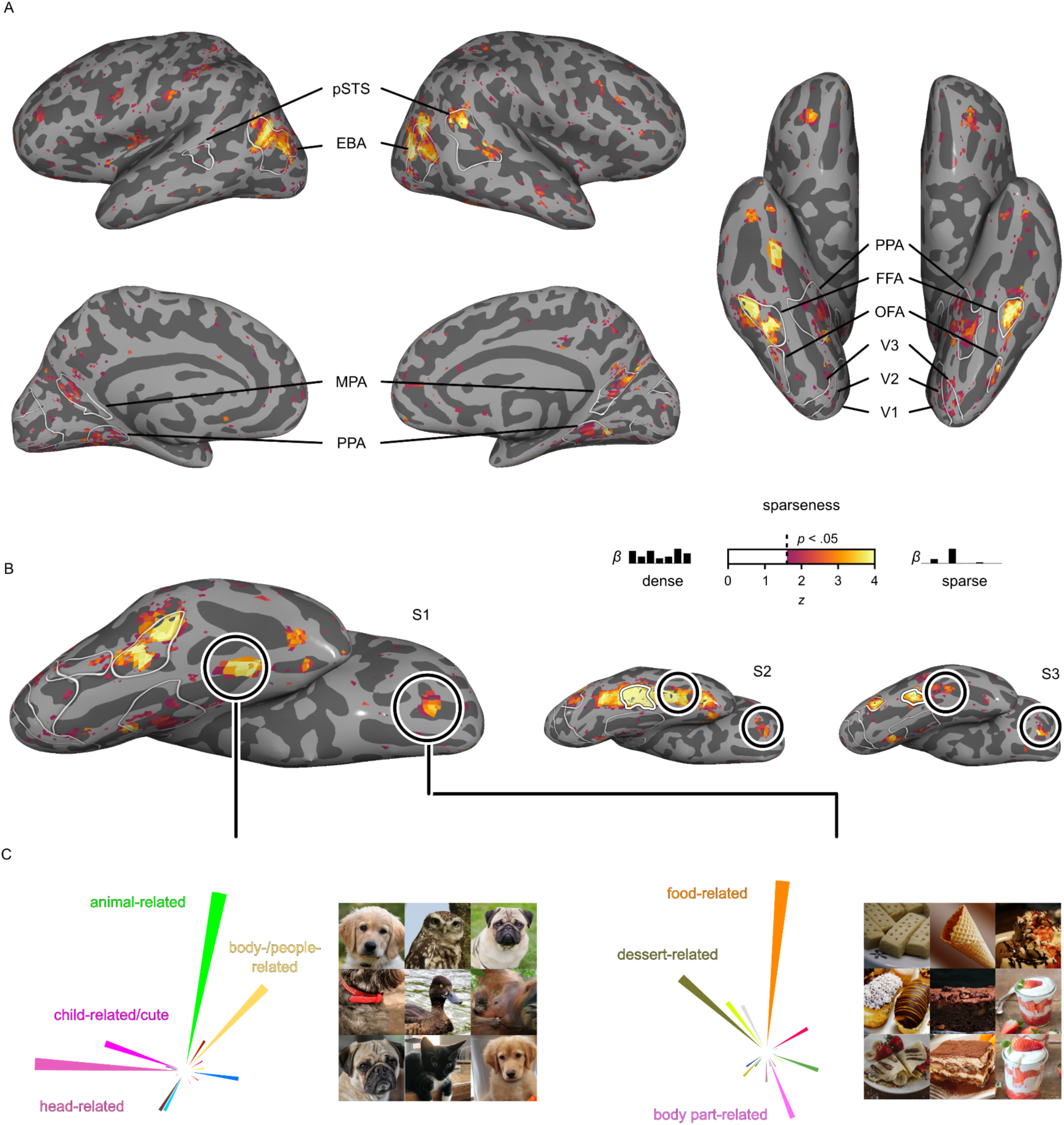
Representational sparseness of behavior-derived object dimensions in object category selective brain regions. A. Inflated cortical surfaces for Subject 1 showing the sparseness over the encoding model weights in each voxel. Colors indicate z-values of sparseness compared to a noise pool of voxels thresholded at p < 0.05 (one-sided, uncorrected). B. Ventral view of the right hemisphere for all three subjects. Round outlines illustrate the location of two explorative, sparsely tuned regions of interest: One in the fusiform gyrus and one in orbitofrontal cortex. C. Functional selectivity of these explorative regions of interest demonstrated by their multidimensional tuning profiles and most representative object images. For copyright reasons, all original images have been replaced with visually similar images. Original images are available upon request.

Beyond the increased sparseness in functionally selective clusters, which had been defined in an independent localizer experiment (Hebart et al., 2023), we explored to what degree we could use sparseness maps for revealing additional, potentially novel functionally selective regions. To this end, we identified two clusters with consistently high sparseness values across subjects (Fig. 5B). One cluster was located in the right hemisphere anterior to anatomically-defined area FG4 (Rosenke et al., 2018) and between functionally-defined FFA and anterior temporal face patch (Rajimehr et al., 2009), with no preferential response to human faces in 2 of 3 subjects in a separate functional localizer. The other cluster was located in orbitofrontal cortex, coinciding with anatomically defined Fo3 between the olfactory and medial orbital sulci (Henssen et al., 2016). Having identified these clusters, we extracted regional tuning profiles and determined the most representative object images for each cluster. Inspection of the tuning profiles in these sparsely tuned regions revealed that their responses were best captured by images of animal faces for the region anterior to FFA and sweet food for orbitofrontal cortex (Fig. 5C). While the results in orbitofrontal cortex are in line with the motivational significance of rewarding foods and food representations in frontal regions (Avery et al., 2023; Jain et al., 2023; Rolls, 2023; Simmons et al., 2005; Small et al., 2007), the selective response to animal faces in the cluster anterior to FFA deserves further study. By identifying regional response selectivity in a data-driven fashion (Lashkari et al., 2010), the results show that sparse tuning can aid in localizing functionally selective brain regions, corroborating the link between representational dimensions and regional selectivity.

### Object dimensions offer a better account of visual cortex responses than categories

If representational dimensions offer a better account of the function of ventral visual cortex than categorization, this would predict that they have superior explanatory power for brain responses to visually-perceived objects in these regions (Downing et al., 2006; Grill-Spector & Weiner, 2014). To compare these accounts formally, we compiled a multidimensional and a categorical model of object responses and compared the amount of shared and unique variance explained by these models (for an exploratory comparison with object shape, see Suppl. Fig. 6 and Suppl. Methods 2). We first constructed a category model by assigning all objects appearing in the presented images into 50 common high-level categories (e.g. "animal", "bird", "body part", "clothing", "food", "fruit", "vehicle") available as part of the THINGS metadata (Stoinski et al., 2023). To account for the known selectivity to faces and body parts, we additionally labeled images in which faces or body parts appeared and included them as two additional categories. Then, for each category, we determined the most diagnostic object dimension. Since some dimensions mapped to multiple categories, this resulted in a model of 30 object dimensions. Based on the 52 categories and the 30 dimensions, we fit two encoding models to the fMRI single trial responses and performed variance partitioning to disentangle the relative contribution of the object category and dimension models to the cross-validated prediction.

The results (Fig. 6) demonstrate that both object dimensions and categories shared a large degree of variance in explaining brain responses, especially in higher-level ventro-temporal and lateral occipital cortices (median = 19%, maximum = 74% shared explained variance) and to a lesser extent in early visual regions (median = 4%, maximum = 19% shared explained variance). This suggests that both models are well suited for predicting responses in the visual system. However, when inspecting the unique variance explained by either model, object dimensions explained a much larger amount of additional variance than object categories (Suppl. Fig. 5). This gain in explained variance was not only evident in higher-level regions (median = 10%, maximum = 35% unique explained variance), where both models performed well, but extended across large parts of visual cortex, including early visual regions (median = 8%, maximum = 35% unique explained variance), suggesting that behavior-derived dimensions captured information not accounted for by categories. Conversely, category membership added little unique explained variance throughout the visual system (median = 1 %, maximum = 11%), reaching larger values in higher-level regions (median = 2%, maximum = 11% unique explained variance). Together, these results indicate that a multidimensional model offers an account with more explanatory value than a category model, supporting the idea that capturing behaviorally-relevant responses in the visual systems requires moving beyond categorization and suggesting object dimensions as a suitable model of encoding the behavioral significance of objects.

**Fig. 6.**
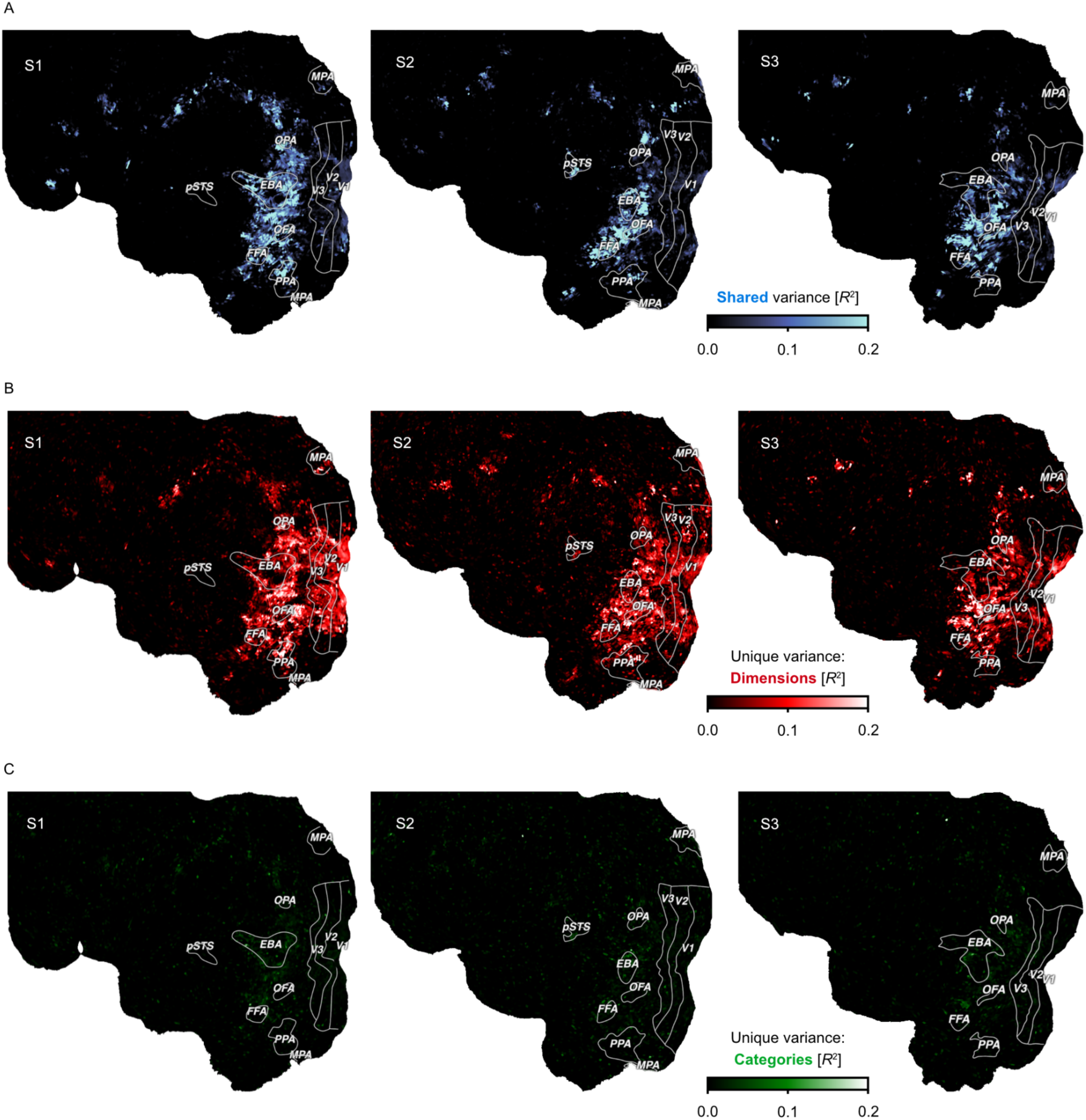
Comparison of a continuous dimensional and a categorical model of object responses. Flat maps show the left hemisphere of each subject. Colors indicate the proportion of explained variance (noise ceiling corrected R^2^) from variance partitioning. A. Shared variance in single-trial fMRI responses explained by both models. B. Variance explained uniquely by a multidimensional model. C. Variance explained uniquely by a model of object categories.

## Discussion

Determining how the human brain represents object properties that inform our broad range of behaviors is crucial for understanding how we make sense of our visual world and act on it in meaningful ways. Here, we identified behavior-derived brain representations by predicting fMRI responses to thousands of object images with 66 interpretable representational dimensions underlying millions of object similarity judgements. The results revealed that this behaviorally-relevant information is mirrored in activity patterns throughout the entire visual processing hierarchy, emphasizing the importance of considering the entire system for identifying the behavioral relevance of visual responses. The diverse image selectivity of different visual brain regions emerged from the multidimensional tuning profiles in this distributed representation. This suggests that behavior-derived dimensions offer a broadly applicable model for understanding the architecture of the visual system in which category-selective regions stand out as a special case of sparse tuning. A direct model comparison confirmed that such a multidimensional account has more explanatory value than a category-centric account.

Much work on the behavioral relevance of object responses in occipitotemporal cortex has focused primarily on a limited number of behavioral goals, such as recognition and high-level categorization (Caramazza & Shelton, 1998; Grill-Spector, 2003; Grill-Spector & Weiner, 2014; Konen et al., 2011; Schiltz et al., 2006; Wada & Yamamoto, 2001). According to this view, high-level visual regions contain representations that abstract from factors non-essential for recognition and categorization, such as position, color, or texture (DiCarlo et al., 2012; DiCarlo & Cox, 2007; Kanwisher, 2010). Our findings provide an alternative perspective onto the nature of cortical object representations that may offer greater explanatory power than this traditional view. By considering a richer representation of objects supporting broader behavioral goals (Bracci & Op de Beeck, 2023), object information is no longer restricted to the commonalities between objects based on how we label them. In this framework, even responses in early visual cortex to images from high-level categories such as food (Jain et al., 2023; Khosla et al., 2022), which would traditionally be disregarded as lower-level confounds based on texture or color, are relevant information supporting the processing of behaviorally-relevant visual inputs. In this perspective, object vision solves the more general problem of providing a rich representation of the visual environment capable of informing a diverse array of behavioral domains (Bracci & Op de Beeck, 2023).

While our results favor a distributed view of object representations, localized response selectivity for ecologically important object stimuli has been replicated consistently, underscoring the significance of specialized clusters. Regional specialization and distributed representations have traditionally been seen as opposing levels of description (Haxby et al., 2001; Op de Beeck et al., 2008). In contrast, our study advances a framework for unifying these perspectives by demonstrating that, compared to other visual regions, category selective clusters exhibit sparse response tuning profiles. This framework treats regional specialization not as an isolated phenomenon, but rather a special case within a more general organizing principle. Thus, it provides a more general view of object representations that acknowledges the significance of regional specialization in the broader context of a multidimensional topography.

One limitation of our study is that we did not identify behavior-derived dimensions specific to each individual participant tested in the MRI. Instead, dimensions were based on a separate population of participants. However, our findings were highly replicable across the three participants for most dimensions, suggesting that these dimensions reflect general organizing principles rather than idiosyncratic effects (Suppl. Fig. 4). Of note, some dimensions did not replicate well (e.g. "feminine (stereotypical)", "hobby-related", or "foot- / walking-related"; Suppl. Fig. 4), which indicates that our fitting procedure does not yield replicable brain activity patterns for any arbitrary dimension. Future work may test the degree to which these results generalize to other dimensions identified through behavior. Additionally, applying our approach to an external fMRI dataset (Suppl. Methods. 1) revealed similarly distributed responses, with highly similar dimension tuning maps, suggesting that our findings generalize to independent participants (Suppl. Fig. 2). Future work could test the extent to which these results generalize to the broader population and how they vary between individuals. Further, despite the broad diversity of objects used in the present study, our work excluded non-object images like text (Hebart et al., 2019). While effects of representational sparseness were less pronounced in scene-selective regions and largely absent in text-selective regions (Puce et al., 1996), our encoding model significantly predicted brain responses in scene-selective regions (Suppl. Fig. 3), indicating validity beyond isolated objects. Future research may extend these insights by exploring additional image classes. Moreover, our use of a pre-trained computational model (Radford et al., 2021) to obtain predicted dimension values might have underestimated the performance of the object embedding in predicting brain responses or may have selectively improved the fit of some dimensions more than that of others. Future studies could test if using empirically measured dimension values for each image would lead to refined dimension maps. Finally, we reported results based on noise-ceiling corrected R^2^ values. While noise-ceiling normalization is common practice when interpreting encoding model results in order to make them more comparable, the degree to which the results would generalize if noise ceilings were much higher could likely only be addressed with much larger, yet similarly broad datasets.

While the behavior-derived dimensions used in this study were highly predictive of perceived similarity judgments and object categorization (Hebart et al., 2020), there are many possible behaviors not captured by this approach. Here, we used representational dimensions underlying similarity judgments to contrast with the category-centric approach. We chose similarity judgments as a common proxy for mental object representations, since they underlie various behavioral goals, including categorization and recognition (Ashby & Perrin, 1988; Edelman, 1998; Hebart et al., 2020; Nosofsky, 1984; Shepard, 1987). Future work could test the extent to which other behaviors or computational approaches carry additional explanatory value (Charest et al., 2014; Devereux et al., 2014; Magri & Konkle, 2019; C. B. Martin et al., 2018; Ritchie & Carlson, 2016). This would also allow establishing the causal relevance of these activity patterns in behavioral readout (Carlson et al., 2014; Ritchie & Carlson, 2016; Singer et al., 2023; Williams et al., 2007).

Given the explanatory power of our dimensional framework, our results may be interpreted as hinting at an alternative explanation of traditional stimulus-driven feature selectivity through the lens of behavioral relevance (Gibson, 1979), where the emergence of feature selectivity may exist because of the potential for efficient behavioral readout. Since the dimensions used in this study likely do not capture all behaviorally-relevant selectivity, our approach does not allow testing this strong assumption. For example, a direct comparison of our embedding with the predictive performance of a Gabor wavelet pyramid model (Kay et al., 2008) or state-of-the-art deep neural network models (Conwell et al., 2023) would neither support nor refute this idea. Future work could specifically target selectivity to individual visual features to determine the degree to which these representations are accessible to behavioral readout and, thus, may alternatively be explained in terms of behavioral relevance, rather than feature selectivity.

In conclusion, our work provides a multidimensional framework that aligns with the rich and diverse behavioral relevance of objects. This approach promises increased explanatory power relative to a category-centric framework and integrates regional specialization within a broader organizing principle, thus offering a promising perspective for understanding how we make sense of our visual world.

## Methods

### THINGS-data

We relied on the openly available THINGS-data collection to investigate the brain representation of every-day objects (Hebart et al., 2023). THINGS-data includes 4.7 million human similarity judgements as well as neural responses measured with functional magnetic resonance imaging (fMRI) to thousands of naturalistic and broadly sampled object images. The collection also includes a representational embedding of core object dimensions learned from the similarity judgments, which predicts unseen human similarity judgements with high accuracy and offers an interpretable account of the mental representation of objects (Hebart et al., 2020, 2023). Here, we used these object dimensions to predict fMRI responses to object images. All data generation and processing methods are described in detail in the original data publication (Hebart et al., 2023) and are only summarized here.

### Participants

The MRI dataset in the THINGS-data collection comprises data from 3 healthy volunteers (2 female, 1 male, mean age: 25.33 years). Participants had normal or corrected-to-normal visual acuity, and were right-handed. The behavioral dataset in the THINGS-data collection was obtained from 12,340 participants through the crowdsourcing platform Amazon Mechanical Turk (6,619 female, 4,400 male, 56 other, 1,065 not reported; mean age: 36.71, std: 11.87, n=5,170 no age reported). Participants provided informed consent in participation and data sharing, and they received financial compensation for taking part in the respective studies. Data acquisition of the THINGS-data collection was approved by the NIH Institutional Review Board (study protocol 93 M-0170, NCT00001360).

### Stimuli

All images were taken from the THINGS database (Hebart et al., 2019). The THINGS database contains 26,107 high-quality, colored images of 1,854 object concepts from a wide range of nameable living and non-living objects, including non-countable substances (e.g. "grass"), faces (e.g. "baby", "boy", "face"), and body parts (e.g. "arm", "leg", "shoulder"). The stimuli presented during functional MRI included 720 object concepts from the THINGS database, with the first 12 examples of each concept selected for a total of 8,640 images. In addition, 100 of the remaining THINGS images were presented repeatedly in each session for the purpose of estimating data reliability.

### Experimental procedure

Participants of the THINGS-fMRI experiment took part in 15-16 scanning sessions, with the first 1-2 sessions serving to acquire individual functional localizers for retinotopic visual areas and category-selective clusters (faces, body parts, scenes, words, and objects). The main fMRI experiment comprised 12 sessions where participants were presented with the 11,040 THINGS images (8,740 unique images, catch trials excluded, 500 ms presentation followed by 4 s of fixation). For details on the procedure of the fMRI and behavioral experiments, please consult the original publication of the datasets (Hebart et al., 2023).

Behavioral similarity judgements in the THINGS-data collection were collected in a triplet odd-one-out study using the online crowdsourcing platform Amazon Mechanical Turk. Participants were presented with three object images side by side and were asked to indicate which object they perceived to be the odd-one-out. Each task comprised 20 odd-one-out trials, and participants could perform as many tasks as they liked.

### MRI data acquisition and preprocessing

Whole-brain functional MRI images were acquired with 2mm isotropic resolution and a repetition time of 1.5s. The MRI data was preprocessed with the standard pipeline fMRIPrep (Esteban et al., 2019) which included slice time correction, head motion correction, susceptibility distortion correction, co-registration between functional and T1-weighted anatomical images, brain tissue segmentation, and cortical surface reconstruction. Additionally, cortical flat maps were manually generated (Gao et al., 2015). Functional MRI data was denoised with a semi-automated procedure based on independent component analysis (ICA) which was developed specifically for the THINGS-fMRI dataset. The retinotopic mapping data and functional localizer data were used to define retinotopic visual regions as well as the category-selective regions used in this study. Image-wise response estimates were obtained by fitting a single-trial model to the fMRI time series of each functional run while accounting for variation in hemodynamic response shape and mitigating overfitting (Allen et al., 2022; Prince et al., 2022; Rokem & Kay, 2020).

### Behavioral embedding

In order to predict the neural response to seen objects, we used a recent, openly available model of representational dimensions underlying human similarity judgements of objects (Hebart et al., 2020). This model was trained to estimate a low-dimensional, sparse, and non-negative embedding predictive of individual trial choices in an odd-one-out task on 1,854 object images. The dimensions of this embedding have been demonstrated to be highly predictive of human similarity judgments while yielding human-interpretable dimensions reflecting both perceptual (e.g. "red", "round") as well as conceptual (e.g. "animal-related") object properties. We used a recent 66-dimensional embedding trained on 4.7 million odd-one-out judgments on triplets of 1,854 object images (Hebart et al., 2023).

While the original embedding was trained on one example image for each of the 1,854 object concepts, it may not account for differences between exemplars of the same object concept. For example, the color of the apple the model was trained on might have been red, while we also presented participants with images of a green apple. This may underestimate the model’s potential to capture variance in brain responses to visually presented object images. To address this, we extended the original embedding by predicting the 66 object dimensions for each individual image in the THINGS database (Hebart et al., 2019). To this end, we used the neural network model CLIP-ViT, which is a multimodal model trained on image-text pairs and which was recently demonstrated to yield excellent prediction of human similarity judgments (Kaniuth et al., 2024; Muttenthaler et al., 2023). For each of the 1,854 object images, we extracted the activity pattern from the final layer of the image encoder. Then, for each of the 66 dimensions, we fitted a ridge regression model to predict dimension values, using cross-validation to determine the regularization hyperparameter. Finally, we applied the learned regression model to activations for all images in the THINGS database. This resulted is a 66-dimensional embedding that captures the mental representation of all 26,107 THINGS images. We used these predicted dimension values to predict fMRI responses to the subset of 8,740 unique images presented in fMRI, which yielded consistent improvements in explained variance for all dimensions (see Suppl. Fig. 1).

### Encoding model

We used a voxel-wise encoding model of the 66-dimensional similarity embedding to predict image-wise fMRI responses in order to test 1) how well the model predicts neural responses in different parts of the visual system and 2) how neural tuning to individual dimensions maps onto the topography of visual cortex.

### Linear regression on fMRI single trial estimates

To test how well the core object dimensions predict brain responses in different parts of the visual system, we fit them to the fMRI single trial response estimates using ordinary least squares regression. While most analyses in this work rely on a more powerful parametric modulation model estimated on time-series data (see below), we used single trial responses for estimating the predictivity of the object dimensions, since this approach does not require extracting the contribution of the parametric modulators for estimating the explained variance of the general linear model. We evaluated the prediction performance of this encoding model in a leave-one-session-out cross-validation, using the average correlation between predicted and observed fMRI responses across folds. Within each cross-validation fold, we also computed a null distribution of correlation values based on 10,000 random permutations of the held-out test data. To assess statistical significance, we obtained voxel-wise *p*-values by comparing the estimated correlation with the generated null-distribution and corrected for multiple comparisons based on a false discovery rate of *p* < 0.01. We computed noise ceiling corrected R^2^ values by dividing the original R^2^ of the model by the noise ceiling estimates, for each voxel separately. These single-trial noise ceilings (Suppl. Fig. 7) were provided with the fMRI dataset and were computed based on estimates of the signal and noise variance obtained based on the variability of responses to repeatedly presented images (Hebart et al., 2023).

### Parametric modulation on fMRI time series

In order to evaluate the contribution of individual object dimensions to the neural response in a given voxel, we used a parametric modulation model on the voxel-wise time series data. In this parametric modulation, a general onset regressor accounts for the average response across all trials, and a set of 66 parametric modulators account for the modulation of the BOLD signal by individual object dimensions. To compile the parametric modulation model, we constructed dimension-specific onset regressors and mean-centered each parametric modulator in order to make them orthogonal to the general onset regressor. We then convolved these regressors with a hemodynamic response function (HRF) to obtain predictors of the BOLD response. To account for variation in the shape of the HRF, we determined the best fitting HRF for each voxel based on a library of 20 HRFs (Allen et al., 2022; Prince et al., 2022). The resulting design matrix was then concatenated and fit to the fMRI time-series data. In order to mitigate overfitting, we regularized the regression weights using fractional ridge regression (Rokem & Kay, 2020). We chose a range of regularization parameters from 0.10 to 0.90 in steps of 0.10 and from 0.90 to 1.00 in steps of 0.01 in order to sample values more densely which reflect less regularization. We determined the best hyperparameter combination (20 HRFs, 26 regularization parameters) for each voxel based on the amount of variance explained in a 12-fold between-session cross-validation. Finally, we fit the model with the best hyperparameter combination per voxel to the entire dataset, yielding 66 statistical maps of regression weights representing the voxel-wise contribution of individual object dimensions in predicting the fMRI signal. The regularization hyperparameter turned out to be small throughout visual cortex (Suppl. Fig. 8), demonstrating that regularization of regression weights had little impact on the absolute size of regression weights. While our analysis was focused on individual subjects, we also estimated the consistency of the tuning maps of individual dimensions across participants. To this end, we used a number of individually-defined regions of interest as anchor points for quantifying similarities and differences between these maps. First, for each dimension separately, we obtained mean beta patterns across these regions, including early visual retinotopic areas (V1-V3 and hV4) as well as face- (FFA, OFA), body- (EBA), and scene-selective (PPA, OPA, MPA) regions. Face-, body-, and scene-selective regions were analyzed separately for each hemisphere to account for potential lateralized effects, and voxels with a noise ceiling smaller than 2% were excluded from the analysis. Finally, to quantify the replicability across participants, we computed the inter-subject correlation based on these mean beta patterns, separately for each dimension (Suppl. Fig. 4).

### Regional tuning profiles and most representative object images

To explore the functional selectivity implied by regional tuning to core object dimensions, we extracted tuning profiles for different visual brain regions and related them to the multidimensional representation of all object images in the THINGS database (Hebart et al., 2019) using a high-throughput approach. First, we extracted the regression weights resulting from the parametric modulation model in different visual brain regions: V1, V2, V3, human V4 (hV4), OFA, FFA, EBA, PPA, MPA, OPA. We then averaged these regional tuning profiles across participants and set negative weights to zero, given that the predicted dimensions reflect non-negative values, as well. We plotted the regional tuning profiles as rose plots to visualize the representation of core object dimensions in these brain regions. In order to explore the regional selectivity for specific object images, we determined the cosine similarity between each regional tuning profile and the model representation of all 26,107 images in the THINGS database. This allowed us to identify those THINGS images that are most representative of the local representational profile in different visual brain regions.

### Representational sparseness

We estimated the sparseness of the representation of core object dimensions based on the regression weights from the parametric modulation model. Given our aim of identifying local clusters of similarly-tuned voxels, we performed spatial smoothing on the regression weight maps (FWHM = 4mm) to increase the spatial signal-to-noise ratio. We then took the vectors representing the 66-dimensional tuning profile for each voxel and removed negative vector elements, mirroring the analysis of the regional tuning profiles. We computed the sparseness of the resulting voxel-wise tuning vectors based on a previously introduced sparseness measure which is based on the normalized relationship between the L-1 and L-2 norm of a vector (Hoyer, 2004):

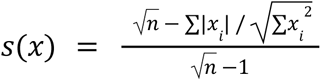

Where *s* indicates the sparseness of the *n*-dimensional input vector *x*. A sparseness value of 1 indicates a perfectly sparse representation where all vector elements except one have the same value. In turn, a value of 0 indicates a perfectly dense representation where all elements have identical values. We computed this sparseness measure over the regression weights in each voxel which yielded a sparseness measure as a single value per voxel. To assess their statistical significance, we first identified the distribution of sparseness values in a noise pool of voxels. This noise pool included voxels where the parametric modulation model predicted the fMRI signal poorly in the cross-validation procedure (*R*^2^ < 0.0001). Since visual inspection of sparseness histograms suggested a log-normal distribution, we log-transformed all sparseness values to convert them to a normal distribution. Finally, we estimated the mean and standard deviation of the sparseness distribution in the noise pool, allowing us to obtain z- and p-values of the sparseness in each voxel.

Based on these results, we explored whether local clusters of representational sparseness are indicative of brain regions with high functional selectivity. To this end, we identified two regional clusters of high sparseness values which were present in all subjects and which had not yet been defined based on the functional localizer experiment (see MRI data preprocessing). Based on visual inspection of the sparseness maps, we defined two regions of interest. The first region of interest was located in the right ventro-temporal cortex, anterior to anatomically-defined area FG4 (Rosenke et al., 2018) and functionally-defined FFA, but posterior to the anterior temporal face patch (Rajimehr et al., 2009). The second region of interest was located in the orbitofrontal cortex. We probed the functional selectivity of these sparsely tuned regions by extracting regional tuning profiles and determining the most representative object images as described in the previous section.

### Variance partitioning of object category vs. dimension based models

The aim of the variance partitioning was to test whether object dimensions or object categories offer a better model of neural responses to object images. To this end, we compiled a multidimensional and categorical model and compared the respective amount of shared and unique variance explained by these models.

We used 50 superordinate object categories provided in the THINGSplus metadata collection to construct a category encoding model (Stoinski et al., 2023) (see Suppl. Methods 3 for a full list). To account for cases where images contained multiple objects (e.g. an image of "ring" might also contain a finger), we used the image annotations in the THINGSplus metadata (Stoinski et al., 2023) and manually matched these annotations to objects in the THINGS database for all images presented in the fMRI experiment. Lastly, we added two more categories by manually identifying images containing human faces or body parts, respectively. We then compiled an encoding model with 52 binary regressors encoding the high-level categories of all respective objects.

Next, we compiled a corresponding encoding model of object dimensions. Note that we predicted that this model would outperform the categorical model in explaining variance in neural responses. To conservatively test this prediction, we biased our analysis in favor of the categorical model by selecting fewer dimensions than categories. To this end, for each category we identified the object dimension with the strongest relationship based on the area under the curve metric (AUC). Since some dimensions are diagnostic for multiple categories (e.g. "animal-related" might be the most diagnostic dimension for both "bird" and "insect"), this resulted in a one-to-many mapping between 30 dimensions and 50 categories (see Suppl. Methods 3 for a full list of selected dimensions).

In order to compare the predictive potential of these two models, we fitted them to the fMRI single trial responses in a voxel-wise linear regression and performed variance partitioning. In order to estimate the uniquely explained variance, we first orthogonalize the target model and the data with respect to the other model (Mumford et al., 2015). This effectively removes the shared variance from both the target model and the data. We then fit the residuals of the target model to the residuals of the data and calculated the coefficient of determination (***R***2) in a 12-fold between-session cross-validation as an estimate of the unique variance explained by the target model. We then estimated the overall variance explained by both models by concatenating the two models, fitting the resulting combined model to the data, and determining the cross-validated ***R***^2^ estimate. Lastly, we computed an estimate of the shared variance explained by the two models by subtracting the uniquely explained variances from the overall explained variance. For visualization purposes, R^2^ values were normalized by the noise ceiling estimates provided with the fMRI dataset (Hebart et al., 2023) (Suppl. Fig. 7). We also visualized the relationship between the performance of both models quantitatively. To that end, we selected voxels with a noise ceiling of greater than 5% in early- (V1-V3) and higher-level (face-, body-, and scene-selective) regions of interest and created scatter plots comparing the variance uniquely explained by the category- and dimensions-based model in these voxels (Suppl. Fig. 5). To summarize the extent of explained variance, we computed median and maximum values for the shared and unique explained variances in these voxels.

## Data availability

The data supporting our analyses were obtained from the publicly available THINGS-fMRI dataset. The fMRI dataset is accessible on OpenNeuro (https://doi.org/10.18112/openneuro.ds004192.v1.0.5) and Figshare (https://doi.org/10.25452/figshare.plus.c.6161151). The object dimensions embedding underlying behavioral similarity judgements which was used to predict the fMRI responses is available at the Open Science Framework repository (https://osf.io/f5rn6/). The higher-level object category labels which were used to construct a categorical model of object responses are part of the THINGSplus metadata and available at the Open Science Framework (https://osf.io/jum2f/). The BOLD5000 fMRI data, including all image stimuli, are openly available on figshare (https://doi.org/10.1184/R1/14456124).

## Code availability

The python code (version 3.7.6) used for data processing, analysis, and visualization in this study is publicly available on GitHub (https://github.com/ViCCo-Group/dimension_encoding).

## Acknowledgements

We thank Philipp Kaniuth for his help with image-wise dimension predictions, Magdalena Holzner for her help with identifying background objects in images and finding copyright free alternative images for publication, and Jacob Prince for sharing cortical flat maps for the BOLD5000 data. This work was supported by a doctoral student fellowship awarded to O.C. by the Max Planck School of Cognition, the Intramural Research Program of the National Institutes of Health (ZIA-MH-002909), under National Institute of Mental Health Clinical Study Protocol 93-M-1070 (NCT00001360), a research group grant by the Max Planck Society awarded to M.N.H., the ERC Starting Grant project COREDIM (ERC-StG-2021-101039712), and the Hessian Ministry of Higher Education, Science, Research and Art (LOEWE Start Professorship to M.N.H. and Excellence Program "The Adaptive Mind"). The funders had no role in study design, data collection and analysis, decision to publish or preparation of the manuscript.

## Author contributions statement

O.C., C.I.B. and M.N.H. conceived the study. O.C. carried out the data analysis and wrote the original draft of the manuscript. C.I.B. and M.N.H. reviewed the manuscript and provided critical feedback. M.N.H. supervised the project.

## Supplementary Information

### Supplementary Methods

#### Supplementary Methods 1: Replication in BOLD5000

In order to test how well our results replicate in an independent dataset, we applied our encoding model of behavior-derived object dimensions to a different openly available large-scale fMRI dataset of visual responses. We chose the BOLD5000 dataset (Chang et al., 2019) because it included object images taken from a different image database, ImageNet (Deng et al., 2009), as well as scene images taken from the MS CoCo database (Lin et al., 2014) that consist of a mixture of individual objects and multiple objects in a scene. This allowed us to evaluate the extent to which our results generalize to different participants and images classes. However, we excluded responses to scene categories from the SUN database (Xiao et al., 2010) since these images were specifically designed to exclude objects. We further excluded participant CSI4 from this analysis since they performed only part of the experiment. The resulting dataset comprised three participants who each saw 3,916 images (1,916 images from ImageNet and 2,000 images from MS CoCo).

We constructed our model of behaviorally-relevant object representation for the BOLD5000 stimuli by obtaining predicted dimension values based on the same procedure which we used for the THINGS images (see Methods section *Behavioral embedding*). We then fit an encoding model to each individual participant where we predict voxel-wise responses to each image based on the 66 object dimensions. Mirroring our main analysis, we fit this model in two different ways, 1) in order to evaluate its overall prediction performance, and 2) to evaluate the contribution to individual dimensions in each voxel. To estimate the prediction accuracy of the entire model, we used a simple cross-validated linear regression analogous to our main analysis (Methods section *Linear Regression on fMRI single trial estimates*). We normalized this prediction accuracy based on noise ceilings which we obtained with the same procedure that was used for the THINGS-fMRI dataset (Hebart et al., 2023). To obtain robust estimates for the contribution of individual dimensions for this prediction, we fit our encoding model again using fractional ridge regression (Rokem & Kay, 2020). We determined the best-fitting regularization parameter for each voxel based on the same parameter grid and cross-validation procedure which we used in our parametric modulation model, except it was applied to discrete response estimates instead of time series data (see Methods section *Parametric modulation on fMRI time series*). Finally, to visualize the spatial extent of the model prediction accuracy and the dimension-wise tuning maps, we visualized the results on cortical flat maps (Gao et al., 2015).

#### Supplementary Methods 2: Variance partitioning of object shape vs. behavior-derived dimensions

In an exploratory analysis, we tested how much variance in neural responses can be explained by object shape relative to the behavior-derived object dimensions. To this end, we used an image-computable model of object shape and compared its explanatory power to our behavior-derived dimensions using variance partitioning.

We first obtained a model of object shape for all stimuli presented in THINGS-fMRI. To this end, we automatically segmented all images using Segment Anything (Kirillov et al., 2023) which we prompted with a CLIP embedding of the object concept labels to further refine results. Images for which the segmentation algorithm failed (n=873) were excluded from this analysis. Next, we used object silhouettes identified through these segmentations as input to an image-computable model of object shape (Morgenstern et al., 2021). This model represents object shape with 22 dimensions, which have been demonstrated to be highly predictive of perceived shape similarity and which reflect latent components underlying more than 100 shape descriptors, such as fourier descriptors, major axis orientation, or shape skeleton (Morgenstern et al., 2021). If a given image contained multiple objects of the same type (e.g. 3 apples), we averaged by averaging over the values of all segmentations for this given image. From these results, we obtained an encoding model of object shape with 22 regressors. We then compared this shape model with our behavior-derived model in a variance partitioning analogous to our comparison with object category (see Methods *Variance partitioning of object category vs. dimension based models*). This allowed us to disentangle the amount of explained variance in neural responses that is uniquely attributable to object shape or the behaviorally-relevant dimensions, or that is shared by both.

#### Supplementary Methods 3: List of object categories and dimensions used in the variance partitioning

For the comparison of object categories and dimensions, we selected 50 superordinate categories and 30 dimensions.

The selected high-level categories included "animal", "bird", "body part", "breakfast food", "candy", "clothing", "clothing accessory", "condiment", "construction equipment", "container", "dessert", "drink", "electronic device", "farm animal", "food", "footwear", "fruit", "furniture", "game", "garden tool", "hardware", "headwear", "home appliance", "home decor", "insect", "jewelry", "kitchen appliance", "kitchen tool", "lighting", "mammal", "medical equipment", "musical instrument", "office supply", "outerwear", "part of car", "plant", "protective clothing", "safety equipment", "school supply", "scientific equipment", "sea animal", "seafood", "sports equipment", "tool", "toy", "vegetable", "vehicle", "watercraft", "weapon", and "women’s clothing".

The selected dimensions comprised "Metallic / artificial", "food-related", "animal-related", "textile", "plant-related", "house-related / furnishing-related", "valuable / precious", "transportation- / movement-related", "electronics / technology", "colorful / playful", "outdoors", "paper-related / flat", "hobby-related / game-related / playing-related", "tools-related / handheld / elongated", "fluid-related / drink-related", "water-related", "weapon-related / war-related / dangerous", "household-related", "feminine (stereotypical)", "body part-related", "music-related / hearing-related / hobby-related / loud", "construction-related / craftsmanship-related / housework-related", "spherical / voluminous", "flying-related / sky-related", "bug-related / non-mammalian / disgusting", "heat-related / fire-related / light-related", "foot-related / walking-related", "head-related", "medicine-related / health-related", and "sweet / dessert-related"

---

**Supplementary Fig. 1.**
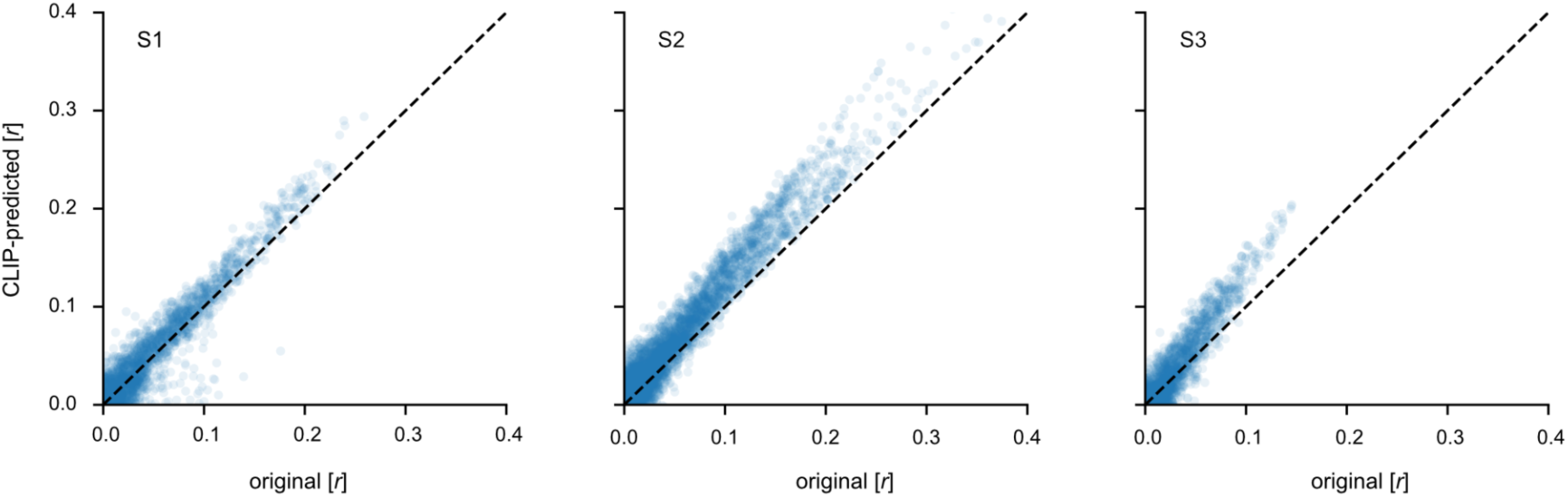
Improvements in fMRI encoding model accuracy after image-wise prediction of object dimensions. Scatter plots show the Pearson correlation between held out and predicted data (12-fold cross-validation) for our voxel-wise fMRI encoding model based on the 66 object dimensions underlying perceived similarity (Hebart et al., 2023). Each sample represents one voxel in a mask of visual cortex (V1-V3, FFA, OFA, pSTS, EBA, PPA, OPA, RSC). The x-axis denotes the prediction performance of the original object embedding based on 1,854 object concepts (Hebart et al., 2023). The y-axis denotes the prediction performance after these original object dimension weights have been predicted for each individual object image presented in fMRI (see Methods section on behavioral model). The dotted line shows equal performance in both cases.

**Supplementary Fig. 2.**
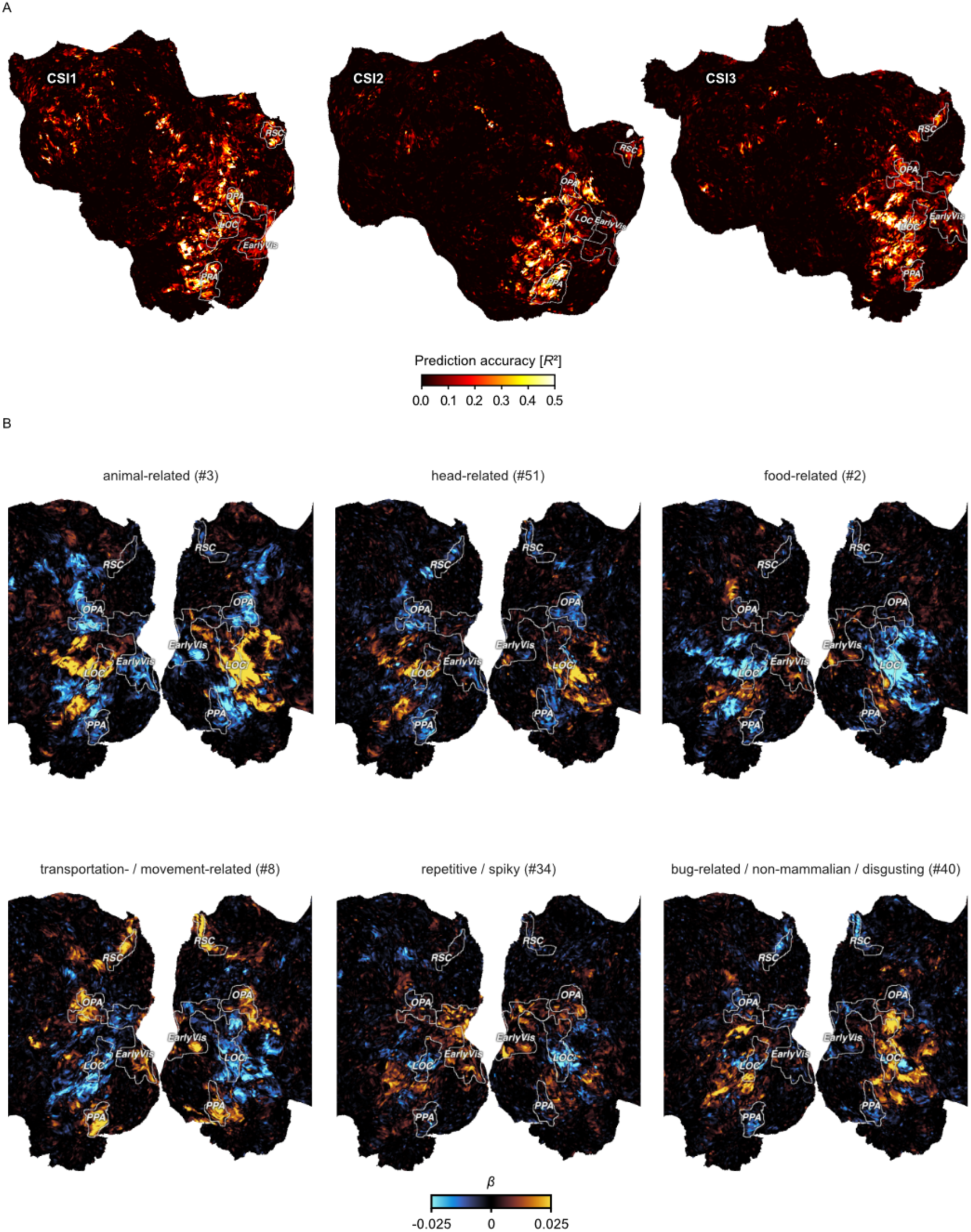
Replication in the BOLD5000 dataset. A. Prediction accuracy of an fMRI encoding model based on the predicted object embedding for BOLD5000 stimuli (noise-corrected *R^2^*). Each column shows the flattened left cortical surface of one subject. The labels “CSI1”, “CSI2”, and “CSI3” correspond to the BOLD5000 subjects. Note that the ROI for early visual cortex is much smaller in BOLD5000 compared to the THINGS-fMRI dataset due to a smaller stimulus presentation (4.6 compared to 10 degree visual angle) and a different procedure for producing flat maps. B. Functional tuning maps for individual object dimensions in example subject CSI3.

**Supplementary Fig. 3.**
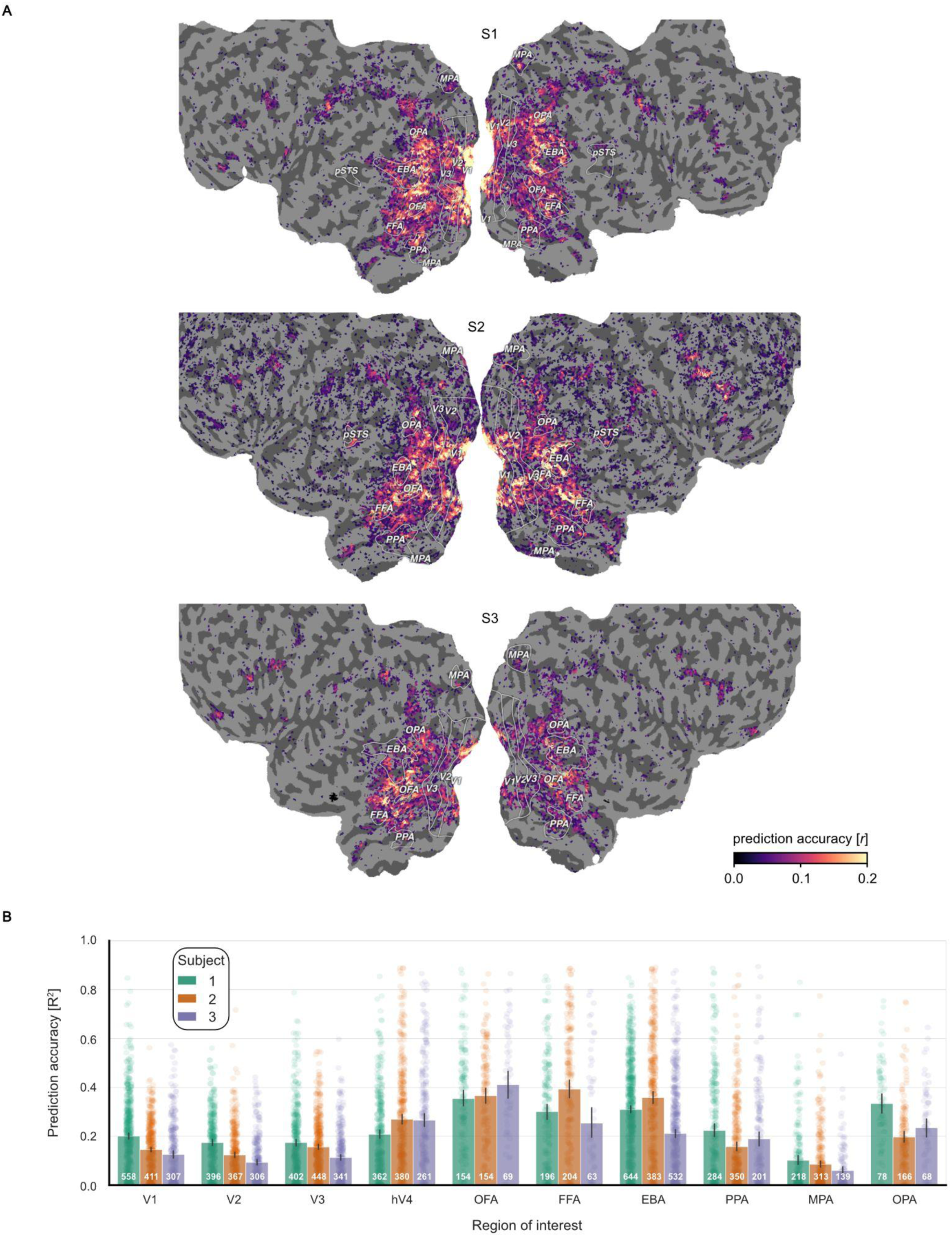
fMRI encoding model prediction accuracy and average accuracy in different ROIs. A. Prediction accuracy in statistically significant voxels (*p*<0.01, FDR-corrected, 12-fold cross-validation, 8,640 training and 820 test samples per fold, 10,000 random permutations per fold). Each row shows flattened cortical surfaces for each subject. Colors indicate Pearson correlation between predicted and held-out data in a between-session 12-fold cross-validation. B. Prediction accuracy in different regions of interest expressed as *R*^2^. Regions of interest include retinotopic areas (V1, V2, V4, hV4) and category-selective clusters (OFA, FFA, EBA, PPA, MPA, OPA). Bars represent the mean value per ROI. Error bars indicate 95% confidence intervals of the mean. Each data point represents one voxel. White annotations indicate the number of voxels per ROI.

**Supplementary Fig. 4.**
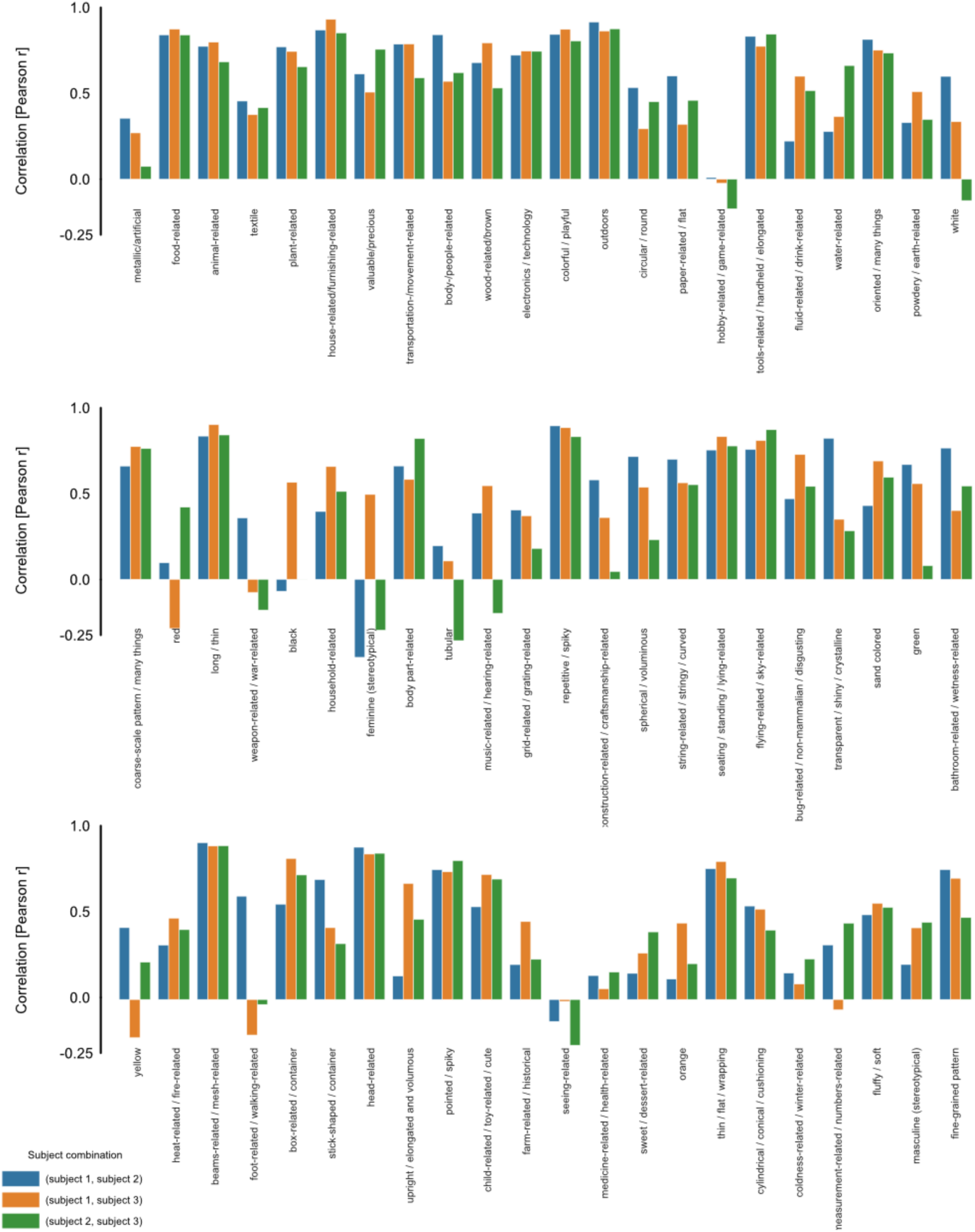
Consistency of average ROI dimension tuning across subjects. Bar heights show the correlation between two participants’ dimension tuning patterns for a given dimension. Tuning patterns were obtained by averaging beta values from the encoding model in 16 ROIs. Bar color indicates the subject pair for which the correlation was computed.

**Supplementary Fig. 5.**
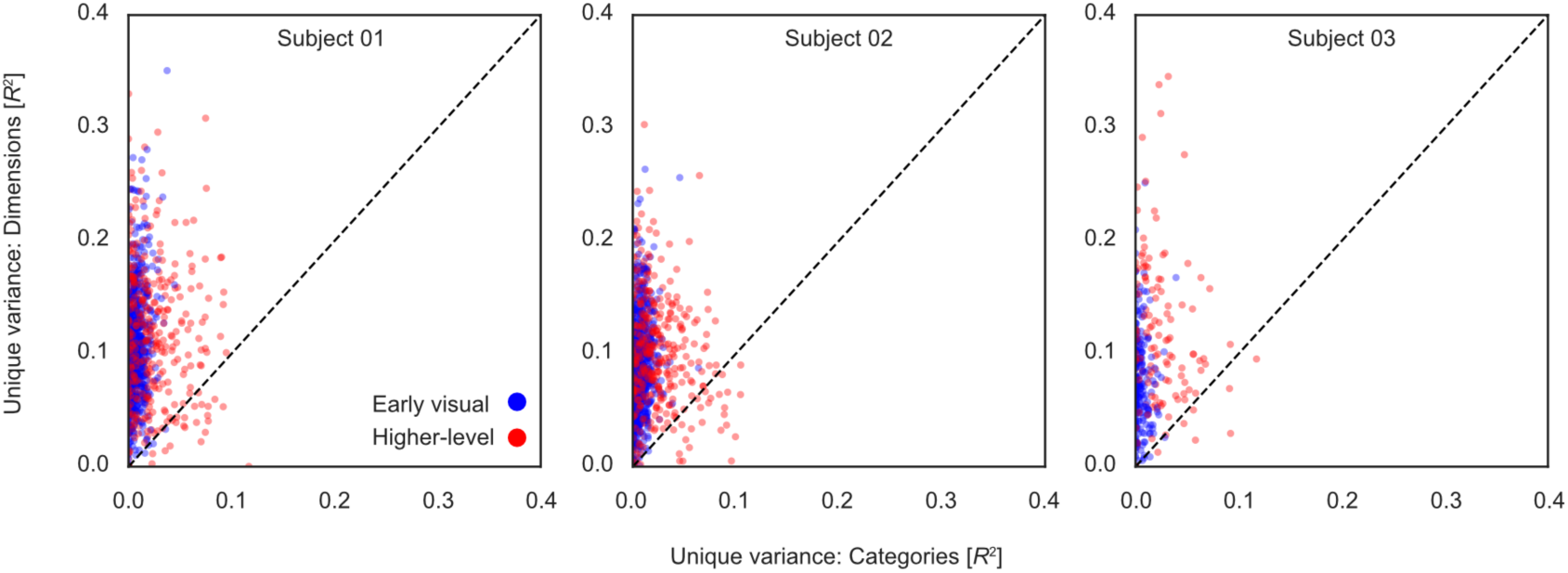
Comparison of variance in neural responses uniquely explained by object category vs. dimensions. Each sample represents one voxel. The x-axis indicates the amount of variance explained by an encoding model of object category, and the y-axis in turn by a model of behavior-derived dimensions. Voxels above the dashed identity line were better explained by the dimensions model. Color indicates whether voxels belong to early-visual (V1-V3) or higher-level (face-, body-, and scene-selective) regions of interest.

**Supplementary Fig. 6.**
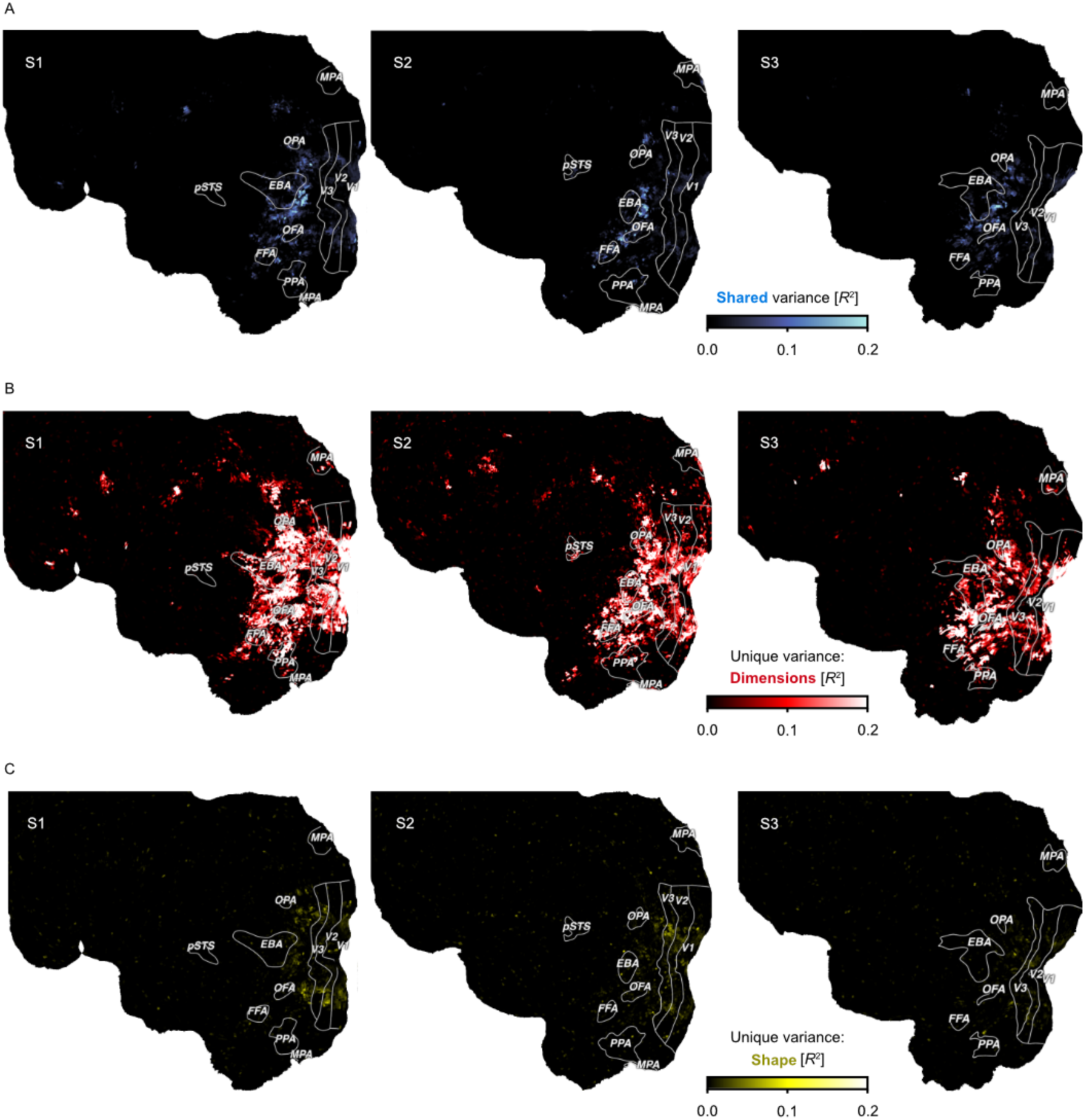
Comparison of a model of object shape and the model of behavior-derived object dimensions. Flat maps show the left hemisphere of each subject. Colors indicate the proportion of explained variance (noise ceiling corrected R^2^) from variance partitioning. A. Shared variance explained by both models. B. Variance explained uniquely by the model of behavioral dimensions. C. Variance explained uniquely by a model of object shape.

**Supplementary Fig. 7.**
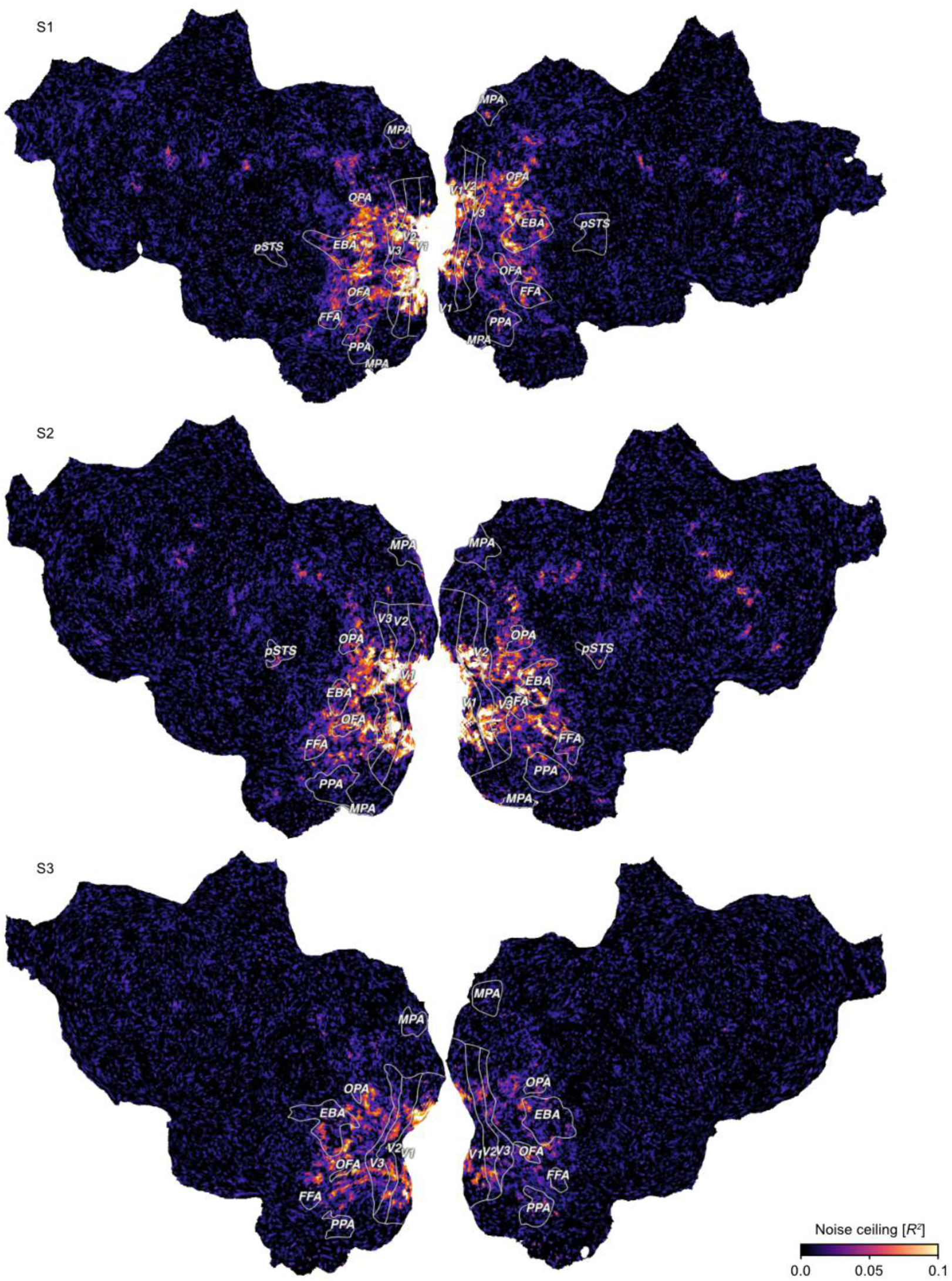
Noise ceiling of single trial responses provided by the THINGS-fMRI dataset. Colors indicate the noise ceiling expressed as the amount of explainable variance in trial-wise fMRI response estimates which was used to normalize the prediction performance of the encoding model.

**Supplementary Fig. 8.**
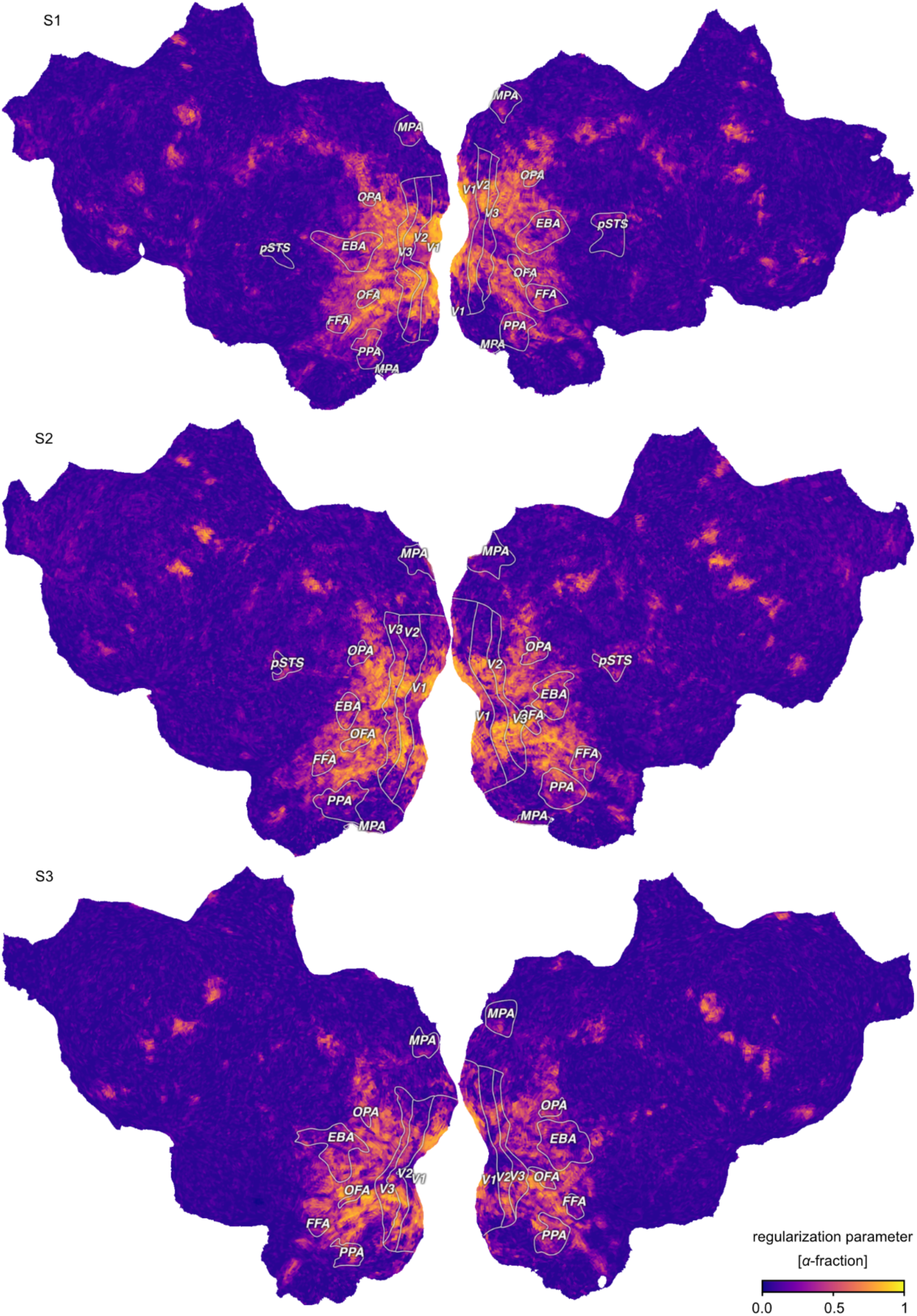
Regularization parameter in the parametric modulation model. Colors indicate voxel-wise *α*-fraction used to regularize the weights in the fractional ridge regression. Larger *α*-fraction reflect a smaller amount of regularization. An *α*-fraction of 1 is equivalent to the ordinary least squares solution. An *α*-fraction of 0 indicates maximum regularization, with all regression weights shrunk to 0.

## References

Allen, E. J., St-Yves, G., Wu, Y., Breedlove, J. L., Prince, J. S., Dowdle, L. T., Nau, M., Caron, B., Pestilli, F., Charest, I., Hutchinson, J. B., Naselaris, T., & Kay, K. (2022). A massive 7T fMRI dataset to bridge cognitive neuroscience and artificial intelligence. Nature Neuroscience, 25(1), 116–126. 10.1038/s41593-021-00962-x

Almeida, J., Fracasso, A., Kristensen, S., Valerio, D., Bergstrom, F., Chakravarthi, R., Tal, Z., & Walbrin, J. (2023). Neural and behavioral signatures of the multidimensionality of manipulable object processing. In bioRxiv. 10.1101/2023.03.29.534804

Andrews, T. J., Clarke, A., Pell, P., & Hartley, T. (2010). Selectivity for low-level features of objects in the human ventral stream. NeuroImage, 49(1), 703–711. 10.1016/j.neuroimage.2009.08.046

Arcaro, M. J., & Livingstone, M. S. (2021). On the relationship between maps and domains in inferotemporal cortex. Nature Reviews. Neuroscience, 22(9), 573–583. 10.1038/s41583-021-00490-4

Ashby, F. G., & Perrin, N. A. (1988). Toward a unified theory of similarity and recognition. Psychological Review, 95(1), 124–150. 10.1037/0033-295X.95.1.124

Avery, J., Carrington, M., Ingeholm, J. E., Simmons, W. K., Hall, K. D., & Martin, A. (2023). Dissociable prefrontal and limbic brain networks represent distinct information about the healthiness and pleasantness of food. In PsyArXiv. 10.31234/osf.io/9qswa

Bao, P., She, L., McGill, M., & Tsao, D. Y. (2020). A map of object space in primate inferotemporal cortex. Nature, 583(7814), 103–108. 10.1038/s41586-020-2350-5

Bracci, S., & Op de Beeck, H. (2016). Dissociations and Associations between Shape and Category Representations in the Two Visual Pathways. The Journal of Neuroscience: The Official Journal of the Society for Neuroscience, 36(2), 432–444. 10.1523/JNEUROSCI.2314-15.2016

Bracci, S., & Op de Beeck, H. P. (2023). Understanding Human Object Vision: A Picture Is Worth a Thousand Representations. Annual Review of Psychology, 74, 113–135. 10.1146/annurev-psych-032720-041031

Caramazza, A., & Shelton, J. R. (1998). Domain-Specific Knowledge Systems in the Brain: The Animate-Inanimate Distinction. Journal of Cognitive Neuroscience, 10(1), 1–34. 10.1162/089892998563752

Carlson, T. A., Ritchie, J. B., Kriegeskorte, N., Durvasula, S., & Ma, J. (2014). Reaction time for object categorization is predicted by representational distance. Journal of Cognitive Neuroscience, 26(1), 132–142. 10.1162/jocn_a_00476

Chang, N., Pyles, J. A., Marcus, A., Gupta, A., Tarr, M. J., & Aminoff, E. M. (2019). BOLD5000, a public fMRI dataset while viewing 5000 visual images. Scientific Data, 6(1), 49. 10.1038/s41597-019-0052-3

Charest, I., Kievit, R. A., Schmitz, T. W., Deca, D., & Kriegeskorte, N. (2014). Unique semantic space in the brain of each beholder predicts perceived similarity. Proceedings of the National Academy of Sciences of the United States of America, 111(40), 14565–14570. 10.1073/pnas.1402594111

Cichy, R. M., Kriegeskorte, N., Jozwik, K. M., van den Bosch, J. J. F., & Charest, I. (2019). The spatiotemporal neural dynamics underlying perceived similarity for real-world objects. NeuroImage, 194, 12–24. 10.1016/j.neuroimage.2019.03.031

Coggan, D. D., Baker, D. H., & Andrews, T. J. (2019). Selectivity for mid-level properties of faces and places in the fusiform face area and parahippocampal place area. The European Journal of Neuroscience, 49(12), 1587–1596. 10.1111/ejn.14327

Coggan, D. D., Liu, W., Baker, D. H., & Andrews, T. J. (2016). Category-selective patterns of neural response in the ventral visual pathway in the absence of categorical information. NeuroImage, 135, 107–114. 10.1016/j.neuroimage.2016.04.060

Coggan, D. D., & Tong, F. (2023). Spikiness and animacy as potential organizing principles of human ventral visual cortex. Cerebral Cortex, 33(13), 8194–8217. 10.1093/cercor/bhad108

Cohen, M. A., Alvarez, G. A., Nakayama, K., & Konkle, T. (2017). Visual search for object categories is predicted by the representational architecture of high-level visual cortex. Journal of Neurophysiology, 117(1), 388–402. 10.1152/jn.00569.2016

Conwell, C., Prince, J. S., Kay, K. N., Alvarez, G. A., & Konkle, T. (2023). What can 1.8 billion regressions tell us about the pressures shaping high-level visual representation in brains and machines? In bioRxiv (p. 2022.03.28.485868). 10.1101/2022.03.28.485868

Cox, D. D. (2014). Do we understand high-level vision? Current Opinion in Neurobiology, 25, 187–193. 10.1016/j.conb.2014.01.016

Deng, J., Dong, W., Socher, R., Li, L.-J., Li, K., & Fei-Fei, L. (2009). ImageNet: A large-scale hierarchical image database. 2009 IEEE Conference on Computer Vision and Pattern Recognition, 248-255. 10.1109/CVPR.2009.5206848

Devereux, B. J., Tyler, L. K., Geertzen, J., & Randall, B. (2014). The Centre for Speech, Language and the Brain (CSLB) concept property norms. Behavior Research Methods, 46(4), 1119–1127. 10.3758/s13428-013-0420-4

DiCarlo, J. J., & Cox, D. D. (2007). Untangling invariant object recognition. Trends in Cognitive Sciences, 11(8), 333–341. 10.1016/j.tics.2007.06.010

DiCarlo, J. J., Zoccolan, D., & Rust, N. C. (2012). How Does the Brain Solve Visual Object Recognition? Neuron, 73(3), 415–434. 10.1016/j.neuron.2012.01.010

Downing, P. E., Chan, A. W.-Y., Peelen, M. V., Dodds, C. M., & Kanwisher, N. (2006). Domain specificity in visual cortex. Cerebral Cortex, 16(10), 1453–1461. 10.1093/cercor/bhj086

Downing, P. E., & Kanwisher, N. (2010). A cortical area specialized for visual processing of the human body. Journal of Vision, 1(3), 341–341. 10.1167/1.3.341

Edelman, S. (1998). Representation is representation of similarities. The Behavioral and Brain Sciences, 21(4), 449–498. 10.1017/s0140525x98001253

Epstein, R. A., Higgins, J. S., & Thompson-Schill, S. L. (2005). Learning places from views: variation in scene processing as a function of experience and navigational ability. Journal of Cognitive Neuroscience, 17(1), 73–83. 10.1162/0898929052879987

Epstein, R. A., & Kanwisher, N. (1998). A cortical representation of the local visual environment. Nature, 392(6676), 598–601. 10.1038/33402

Esteban, O., Markiewicz, C. J., Blair, R. W., Moodie, C. A., Isik, A. I., Erramuzpe, A., Kent, J. D., Goncalves, M., DuPre, E., Snyder, M., Oya, H., Ghosh, S. S., Wright, J., Durnez, J., Poldrack, R. A., & Gorgolewski, K. J. (2019). fMRIPrep: a robust preprocessing pipeline for functional MRI. Nature Methods, 16(1), 111–116. 10.1038/s41592-018-0235-4

Gao, J. S., Huth, A. G., Lescroart, M. D., & Gallant, J. L. (2015). Pycortex: an interactive surface visualizer for fMRI. Frontiers in Neuroinformatics, 9, 23. 10.3389/fninf.2015.00023

Gauthier, I., Skudlarski, P., Gore, J. C., & Anderson, A. W. (2000). Expertise for cars and birds recruits brain areas involved in face recognition. Nature Neuroscience, 3(2), 191–197. 10.1038/72140

Gibson, J. J. (1979). The ecological approach to visual perception. Houghton, Mifflin and Company.

Goodale, M. A., & Milner, A. D. (1992). Separate visual pathways for perception and action. Trends in Neurosciences, 15(1), 20–25. 10.1016/0166-2236(92)90344-8

Grill-Spector, K. (2003). The neural basis of object perception. Current Opinion in Neurobiology, 13(2), 159–166. 10.1016/S0959-4388(03)00040-0

Grill-Spector, K., & Weiner, K. S. (2014). The functional architecture of the ventral temporal cortex and its role in categorization. Nature Reviews. Neuroscience, 15(8), 536–548. 10.1038/nrn3747

Groen, I. I. A., Silson, E. H., & Baker, C. I. (2017). Contributions of low- and high-level properties to neural processing of visual scenes in the human brain. Philosophical Transactions of the Royal Society of London. Series B, Biological Sciences, 372(1714), 20160102. 10.1098/rstb.2016.0102

Hasson, U., Harel, M., Levy, I., & Malach, R. (2003). Large-scale mirror-symmetry organization of human occipito-temporal object areas. Neuron, 37(6), 1027–1041. 10.1016/s0896-6273(03)00144-2

Haxby, J. V., Gobbini, M. I., Furey, M. L., Ishai, A., Schouten, J. L., & Pietrini, P. (2001). Distributed and overlapping representations of faces and objects in ventral temporal cortex. Science, 293(5539), 2425–2430. 10.1126/science.1063736

Hebart, M. N., Contier, O., Teichmann, L., Rockter, A. H., Zheng, C. Y., Kidder, A., Corriveau, A., Vaziri-Pashkam, M., & Baker, C. I. (2023). THINGS-data, a multimodal collection of large-scale datasets for investigating object representations in human brain and behavior. eLife, 12, e82580. 10.7554/eLife.82580

Hebart, M. N., Dickter, A. H., Kidder, A., Kwok, W. Y., Corriveau, A., Van Wicklin, C., & Baker, C. I. (2019). THINGS: A database of 1,854 object concepts and more than 26,000 naturalistic object images. PloS One, 14(10), e0223792. 10.1371/journal.pone.0223792

Hebart, M. N., Zheng, C. Y., Pereira, F., & Baker, C. I. (2020). Revealing the multidimensional mental representations of natural objects underlying human similarity judgements. Nature Human Behaviour, 4(11), 1173–1185. 10.1038/s41562-020-00951-3

He, C., Hung, S.-C., & Cheung, O. S. (2020). Roles of Category, Shape, and Spatial Frequency in Shaping Animal and Tool Selectivity in the Occipitotemporal Cortex. The Journal of Neuroscience: The Official Journal of the Society for Neuroscience, 40(29), 5644–5657. 10.1523/JNEUROSCI.3064-19.2020

Henssen, A., Zilles, K., Palomero-Gallagher, N., Schleicher, A., Mohlberg, H., Gerboga, F., Eickhoff, S. B., Bludau, S., & Amunts, K. (2016). Cytoarchitecture and probability maps of the human medial orbitofrontal cortex. Cortex, 75, 87–112. 10.1016/j.cortex.2015.11.006

Hoyer, P. O. (2004). Non-negative Matrix Factorization with Sparseness Constraints. Journal of Machine Learning Research: JMLR, 5, 1457–1469.

Hubel, D. H., & Wiesel, T. N. (1962). Receptive fields, binocular interaction and functional architecture in the cat’s visual cortex. The Journal of Physiology, 160(1), 106–154. 10.1113/jphysiol.1962.sp006837

Hubel, D. H., & Wiesel, T. N. (1968). Receptive fields and functional architecture of monkey striate cortex. The Journal of Physiology, 195(1), 215–243. 10.1113/jphysiol.1968.sp008455

Hung, C. P., Kreiman, G., Poggio, T., & DiCarlo, J. J. (2005). Fast Readout of Object Identity from Macaque Inferior Temporal Cortex. Science, 310(5749), 863–866. 10.1126/science.1117593

Huth, A. G., Nishimoto, S., Vu, A. T., & Gallant, J. L. (2012). A Continuous Semantic Space Describes the Representation of Thousands of Object and Action Categories across the Human Brain. Neuron, 76(6), 1210–1224. 10.1016/j.neuron.2012.10.014

Jain, N., Wang, A., Henderson, M. M., Lin, R., Prince, J. S., Tarr, M. J., & Wehbe, L. (2023). Selectivity for food in human ventral visual cortex. Communications Biology, 6(1), 175. 10.1038/s42003-023-04546-2

Kaniuth, P., Mahner, F. P., Perkuhn, J., & Hebart, M. N. (2024). A high-throughput approach for the efficient prediction of perceived similarity of natural objects.

Kanwisher, N. (2010). Functional specificity in the human brain: a window into the functional architecture of the mind. Proceedings of the National Academy of Sciences of the United States of America, 107(25), 11163–11170. 10.1073/pnas.1005062107

Kanwisher, N., & Barton, J. J. S. (2011). The functional architecture of the face system: Integrating evidence from fMRI and patient studies. In The Oxford handbook of face perception. Oxford University Press Oxford, UK.

Kanwisher, N., McDermott, J., & Chun, M. M. (1997). The Fusiform Face Area: A Module in Human Extrastriate Cortex Specialized for Face Perception. The Journal of Neuroscience, 17(11), 4302–4311. 10.1523/JNEUROSCI.17-11-04302.1997

Kanwisher, N., & Yovel, G. (2006). The fusiform face area: a cortical region specialized for the perception of faces. Philosophical Transactions of the Royal Society of London. Series B, Biological Sciences, 361(1476), 2109–2128. 10.1098/rstb.2006.1934

Kastner, S., De Weerd, P., Pinsk, M. A., Elizondo, M. I., Desimone, R., & Ungerleider, L. G. (2001). Modulation of sensory suppression: implications for receptive field sizes in the human visual cortex. Journal of Neurophysiology, 86(3), 1398–1411. 10.1152/jn.2001.86.3.1398

Kay, K. N., Naselaris, T., Prenger, R. J., & Gallant, J. L. (2008). Identifying natural images from human brain activity. Nature, 452(7185), 352–355. 10.1038/nature06713

Khosla, M., Ratan Murty, N. A., & Kanwisher, N. (2022). A highly selective response to food in human visual cortex revealed by hypothesis-free voxel decomposition. Current Biology, 32(19), 4159–4171.e9. 10.1016/j.cub.2022.08.009

Kirillov, A., Mintun, E., Ravi, N., Mao, H., Rolland, C., Gustafson, L., Xiao, T., Whitehead, S., Berg, A. C., Lo, W.-Y., Dollar, P., & Girshick, R. (2023). Segment Anything. In *arXiv [cs.CV]*. arXiv. http://arxiv.org/abs/2304.02643

Konen, C. S., Behrmann, M., Nishimura, M., & Kastner, S. (2011). The functional neuroanatomy of object agnosia: a case study. Neuron, 71(1), 49–60. 10.1016/j.neuron.2011.05.030

Konkle, T., & Caramazza, A. (2013). Tripartite Organization of the Ventral Stream by Animacy and Object Size. Journal of Neuroscience, 33(25), 10235–10242. 10.1523/JNEUROSCI.0983-13.2013

Konkle, T., & Oliva, A. (2012). A real-world size organization of object responses in occipitotemporal cortex. Neuron, 74(6), 1114–1124. 10.1016/j.neuron.2012.04.036

Krakauer, J. W., Ghazanfar, A. A., Gomez-Marin, A., MacIver, M. A., & Poeppel, D. (2017). Neuroscience Needs Behavior: Correcting a Reductionist Bias. Neuron, 93(3), 480–490. 10.1016/j.neuron.2016.12.041

Kravitz, D. J., Saleem, K. S., Baker, C. I., Ungerleider, L. G., & Mishkin, M. (2013). The ventral visual pathway: an expanded neural framework for the processing of object quality. Trends in Cognitive Sciences, 17(1), 26–49. 10.1016/j.tics.2012.10.011

Kriegeskorte, N. (2009). Relating Population-Code Representations between Man, Monkey, and Computational Models. Frontiers in Neuroscience, 3(3), 363–373. 10.3389/neuro.01.035.2009

Kriegeskorte, N., Mur, M., Ruff, D. A., Kiani, R., Bodurka, J., Esteky, H., Tanaka, K., & Bandettini, P. A. (2008). Matching categorical object representations in inferior temporal cortex of man and monkey. Neuron, 60(6), 1126–1141. 10.1016/j.neuron.2008.10.043

Lashkari, D., Vul, E., Kanwisher, N., & Golland, P. (2010). Discovering structure in the space of fMRI selectivity profiles. NeuroImage, 50(3), 1085–1098. 10.1016/j.neuroimage.2009.12.106

Lin, T.-Y., Maire, M., Belongie, S., Hays, J., Perona, P., Ramanan, D., Dollar, P., & Zitnick, C. L. (2014). Microsoft COCO: Common Objects in Context. Computer Vision – ECCV 2014, 740–755. 10.1007/978-3-319-10602-1_48

Livingstone, M. S., & Hubel, D. H. (1984). Anatomy and physiology of a color system in the primate visual cortex. The Journal of Neuroscience, 4(1), 309–356. 10.1523/JNEUROSCI.04-01-00309.1984

Magri, C., & Konkle, T. (2019). Comparing facets of behavioral object representation: implicit perceptual similarity matches brains and models. In 2019 Conference on Cognitive Computational Neuroscience. 10.32470/ccn.2019.1395-0

Mahon, B. Z., & Caramazza, A. (2011). What drives the organization of object knowledge in the brain? Trends in Cognitive Sciences, 15(3), 97–103. 10.1016/j.tics.2011.01.004

Marr, D. (2010). Vision: A Computational Investigation into the Human Representation and Processing of Visual Information. MIT Press. https://play.google.com/store/books/details?id=D8XxCwAAQBAJ

Martin, A. (2007). The representation of object concepts in the brain. Annual Review of Psychology, 58, 25–45. 10.1146/annurev.psych.57.102904.190143

Martin, A., & Weisberg, J. (2003). Neural foundations for understanding social and mechanical concepts. Cognitive Neuropsychology, 20(3-6), 575–587. 10.1080/02643290342000005

Martin, A., Wiggs, C. L., Ungerleider, L. G., & Haxby, J. V. (1996). Neural correlates of category-specific knowledge. Nature, 379(6566), 649–652. 10.1038/379649a0

Martin, C. B., Douglas, D., Newsome, R. N., Man, L. L., & Barense, M. D. (2018). Integrative and distinctive coding of visual and conceptual object features in the ventral visual stream. eLife, 7. 10.7554/eLife.31873

Mishkin, M., & Ungerleider, L. G. (1982). Contribution of striate inputs to the visuospatial functions of parieto-preoccipital cortex in monkeys. Behavioural Brain Research, 6(1), 57–77. 10.1016/0166-4328(82)90081-X

Morgenstern, Y., Hartmann, F., Schmidt, F., Tiedemann, H., Prokott, E., Maiello, G., & Fleming, R. W. (2021). An image-computable model of human visual shape similarity. PLoS Computational Biology, 17(6), e1008981. 10.1371/journal.pcbi.1008981

Moro, V., Urgesi, C., Pernigo, S., Lanteri, P., Pazzaglia, M., & Aglioti, S. M. (2008). The neural basis of body form and body action agnosia. Neuron, 60(2), 235–246. 10.1016/j.neuron.2008.09.022

Mumford, J. A., Poline, J.-B., & Poldrack, R. A. (2015). Orthogonalization of regressors in FMRI models. PloS One, 10(4), e0126255. 10.1371/journal.pone.0126255

Mur, M., Meys, M., Bodurka, J., Goebel, R., Bandettini, P. A., & Kriegeskorte, N. (2013). Human Object-Similarity Judgments Reflect and Transcend the Primate-IT Object Representation. Frontiers in Psychology, 4, 128. 10.3389/fpsyg.2013.00128

Muttenthaler, L., & Hebart, M. N. (2021). THINGSvision: A Python Toolbox for Streamlining the Extraction of Activations From Deep Neural Networks. Frontiers in Neuroinformatics, 15, 679838. 10.3389/fninf.2021.679838

Muttenthaler, L., Linhardt, L., Dippel, J., Vandermeulen, R. A., Hermann, K., Lampinen, A., & Kornblith, S. (2023). Improving neural network representations using human similarity judgments. In A. Oh, T. Neumann, A. Globerson, K. Saenko, M. Hardt, & S. Levine (Eds.), Advances in Neural Information Processing Systems (Vol. 36, pp. 50978-51007). Curran Associates, Inc. https://proceedings.neurips.cc/paper_files/paper/2023/file/9febda1c8344cc5f2d51713964864e93-Paper-Conference.pdf

Naselaris, T., Allen, E., & Kay, K. (2021). Extensive sampling for complete models of individual brains. Current Opinion in Behavioral Sciences, 40, 45–51. 10.1016/j.cobeha.2020.12.008

Nasr, S., Echavarria, C. E., & Tootell, R. B. H. (2014). Thinking outside the box: rectilinear shapes selectively activate scene-selective cortex. The Journal of Neuroscience, 34(20), 6721–6735. 10.1523/JNEUROSCI.4802-13.2014

Nasr, S., & Tootell, R. B. H. (2012). A cardinal orientation bias in scene-selective visual cortex. The Journal of Neuroscience, 32(43), 14921–14926. 10.1523/JNEUROSCI.2036-12.2012

Nosofsky, R. M. (1984). Choice, similarity, and the context theory of classification. Journal of Experimental Psychology. Learning, Memory, and Cognition, 10(1), 104–114. 10.1037//0278-7393.10.1.104

O’Craven, K. M., & Kanwisher, N. (2000). Mental imagery of faces and places activates corresponding stiimulus-specific brain regions. Journal of Cognitive Neuroscience, 12(6), 1013–1023. 10.1162/08989290051137549

Op de Beeck, H. P., Haushofer, J., & Kanwisher, N. G. (2008). Interpreting fMRI data: maps, modules and dimensions. Nature Reviews. Neuroscience, 9(2), 123–135. 10.1038/nrn2314

Peelen, M. V., & Downing, P. E. (2017). Category selectivity in human visual cortex: Beyond visual object recognition. Neuropsychologia, 105(XXX), 177–183. 10.1016/j.neuropsychologia.2017.03.033

Pennock, I. M. L., Racey, C., Allen, E. J., Wu, Y., Naselaris, T., Kay, K. N., Franklin, A., & Bosten, J. M. (2023). Color-biased regions in the ventral visual pathway are food selective. Current Biology: CB, 33(1), 134–146.e4. 10.1016/j.cub.2022.11.063

Prince, J. S., Charest, I., Kurzawski, J. W., Pyles, J. A., Tarr, M. J., & Kay, K. N. (2022). Improving the accuracy of single-trial fMRI response estimates using GLMsingle. eLife, 11. 10.7554/eLife.77599

Puce, A., Allison, T., Asgari, M., Gore, J. C., & McCarthy, G. (1996). Differential sensitivity of human visual cortex to faces, letterstrings, and textures: a functional magnetic resonance imaging study. The Journal of Neuroscience, 16(16), 5205–5215. https://www.ncbi.nlm.nih.gov/pubmed/8756449

Radford, A., Kim, J. W., Hallacy, C., Ramesh, A., Goh, G., Agarwal, S., Sastry, G., Askell, A., Mishkin, P., Clark, J., Krueger, G., & Sutskever, I. (2021). Learning Transferable Visual Models From Natural Language Supervision. In M. Meila & T. Zhang (Eds.), Proceedings of the 38th International Conference on Machine Learning (Vol. 139, pp. 8748-8763). PMLR. https://proceedings.mlr.press/v139/radford21a.html

Rajimehr, R., Young, J. C., & Tootell, R. B. H. (2009). An anterior temporal face patch in human cortex, predicted by macaque maps. Proceedings of the National Academy of Sciences, 106(6), 1995–2000. 10.1073/pnas.0807304106

Rice, G. E., Watson, D. M., Hartley, T., & Andrews, T. J. (2014). Low-level image properties of visual objects predict patterns of neural response across category-selective regions of the ventral visual pathway. The Journal of Neuroscience, 34(26), 8837–8844. 10.1523/JNEUROSCI.5265-13.2014

Ritchie, J. B., & Carlson, T. A. (2016). Neural Decoding and "Inner" Psychophysics: A Distance-to-Bound Approach for Linking Mind, Brain, and Behavior. Frontiers in Neuroscience, 10, 190. 10.3389/fnins.2016.00190

Ritchie, J. B., Tovar, D. A., & Carlson, T. A. (2015). Emerging Object Representations in the Visual System Predict Reaction Times for Categorization. PLoS Computational Biology, 11(6), e1004316. 10.1371/journal.pcbi.1004316

Rokem, A., & Kay, K. (2020). Fractional ridge regression: a fast, interpretable reparameterization of ridge regression. GigaScience, 9(12). 10.1093/gigascience/giaa133

Rolls, E. T. (2023). The orbitofrontal cortex, food reward, body weight and obesity. Social Cognitive and Affective Neuroscience, 18(1). 10.1093/scan/nsab044

Rosenke, M., Weiner, K. S., Barnett, M. A., Zilles, K., Amunts, K., Goebel, R., & Grill-Spector, K. (2018). A cross-validated cytoarchitectonic atlas of the human ventral visual stream. NeuroImage, 170, 257–270. 10.1016/j.neuroimage.2017.02.040

Schiltz, C., Sorger, B., Caldara, R., Ahmed, F., Mayer, E., Goebel, R., & Rossion, B. (2006). Impaired face discrimination in acquired prosopagnosia is associated with abnormal response to individual faces in the right middle fusiform gyrus. Cerebral Cortex, 16(4), 574–586. 10.1093/cercor/bhj005

Shepard, R. N. (1987). Toward a universal law of generalization for psychological science. Science, 237(4820), 1317–1323. 10.1126/science.3629243

Silson, E. H., Steel, A. D., & Baker, C. I. (2016). Scene-Selectivity and Retinotopy in Medial Parietal Cortex. Frontiers in Human Neuroscience, 10, 412. 10.3389/fnhum.2016.00412

Simmons, W. K., Martin, A., & Barsalou, L. W. (2005). Pictures of appetizing foods activate gustatory cortices for taste and reward. Cerebral Cortex, 15(10), 1602–1608. 10.1093/cercor/bhi038

Singer, J. J. D., Karapetian, A., Hebart, M. N., & Cichy, R. M. (2023). The link between visual representations and behavior in human scene perception. In bioRxiv (p. 2023.08.17.553708). 10.1101/2023.08.17.553708

Small, D. M., Bender, G., Veldhuizen, M. G., Rudenga, K., Nachtigal, D., & Felsted, J. (2007). The role of the human orbitofrontal cortex in taste and flavor processing. Annals of the New York Academy of Sciences, 1121, 136–151. 10.1196/annals.1401.002

Smith, A. T., Singh, K. D., Williams, A. L., & Greenlee, M. W. (2001). Estimating receptive field size from fMRI data in human striate and extrastriate visual cortex. Cerebral Cortex, 11(12), 1182–1190. 10.1093/cercor/11.12.1182

Stoinski, L. M., Perkuhn, J., & Hebart, M. N. (2023). THINGSplus: New norms and metadata for the THINGS database of 1854 object concepts and 26,107 natural object images. Behavior Research Methods. 10.3758/s13428-023-02110-8

Tootell, R. B., Mendola, J. D., Hadjikhani, N. K., Ledden, P. J., Liu, A. K., Reppas, J. B., Sereno, M. I., & Dale, A. M. (1997). Functional analysis of V3A and related areas in human visual cortex. The Journal of Neuroscience, 17(18), 7060–7078. 10.1523/JNEUROSCI.17-18-07060.1997

Wada, Y., & Yamamoto, T. (2001). Selective impairment of facial recognition due to a haematoma restricted to the right fusiform and lateral occipital region. Journal of Neurology, Neurosurgery, and Psychiatry, 71(2), 254–257. 10.1136/jnnp.71.2.254

Wang, A. Y., Kay, K., Naselaris, T., Tarr, M. J., & Wehbe, L. (2023). Better models of human high-level visual cortex emerge from natural language supervision with a large and diverse dataset. Nature Machine Intelligence, 1-12. 10.1038/s42256-023-00753-y

Williams, M. A., Dang, S., & Kanwisher, N. G. (2007). Only some spatial patterns of fMRI response are read out in task performance. Nature Neuroscience, 10(6), 685–686. 10.1038/nn1900

Xiao, J., Hays, J., Ehinger, K. A., Oliva, A., & Torralba, A. (2010). SUN database: Large-scale scene recognition from abbey to zoo. 2010 IEEE Computer Society Conference on Computer Vision and Pattern Recognition, 3485-3492. 10.1109/CVPR.2010.5539970

Yargholi, E., & Op de Beeck, H. (2023). Category Trumps Shape as an Organizational Principle of Object Space in the Human Occipitotemporal Cortex. The Journal of Neuroscience, 43(16), 2960–2972. 10.1523/JNEUROSCI.2179-22.2023

Zheng, C. Y., Pereira, F., Baker, C. I., & Hebart, M. N. (2019). Revealing interpretable object representations from human behavior (pp. 1–16). arXiv. 10.48550/arXiv.1901.02915

