## Extended Data Figures for "Distributed representations of behavior-derived object dimensions in the human visual system"

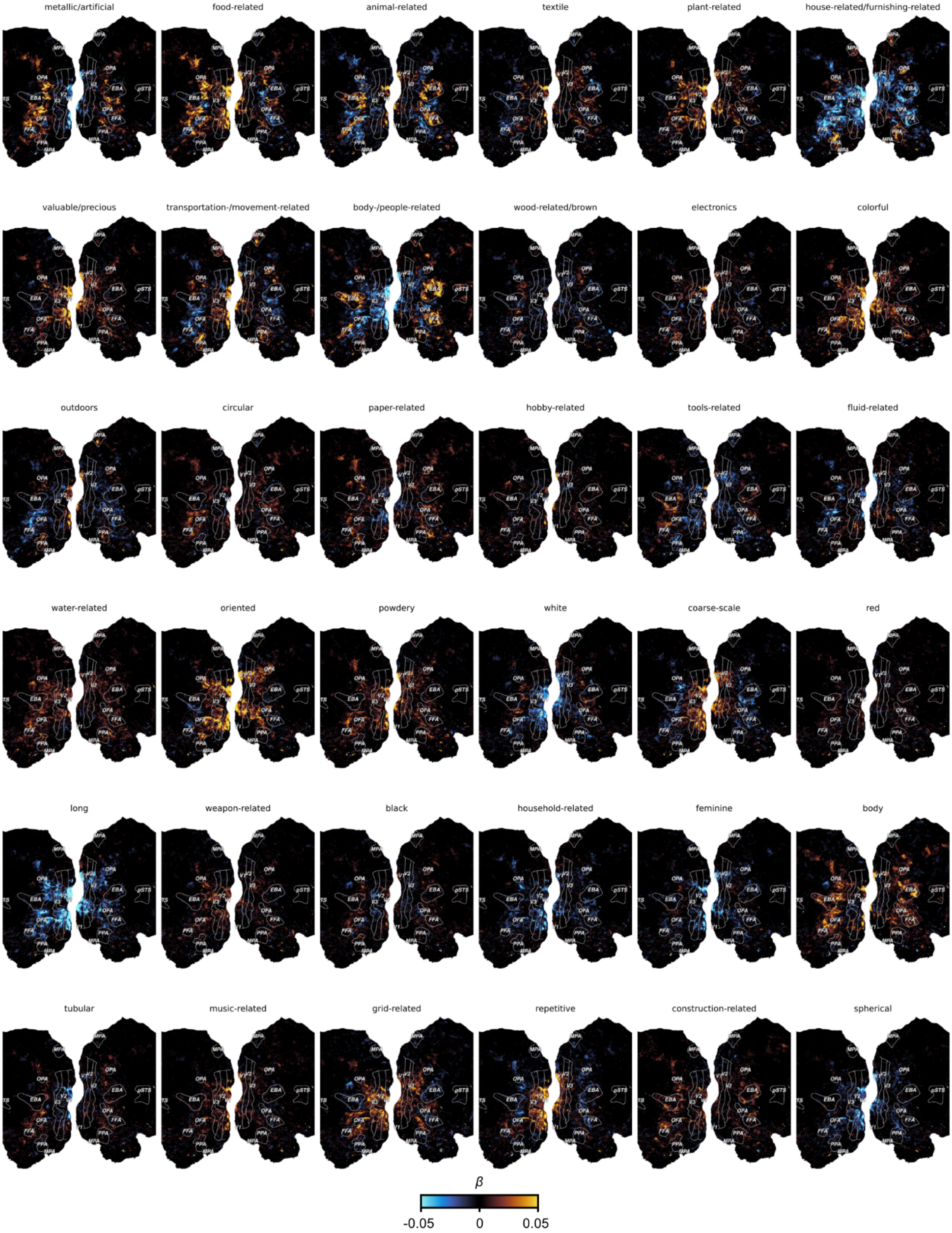


Extended Data Figure 1. **Dimension tuning maps 1-36 for Subject 1**. Colors indicate regression weights for each dimension predictor from the parametric modulation encoding model. Labels indicate regions of interest on the cortex: V1-V3: primary - tertiary visual cortex, OFA: occipital face area, FFA: fusiform face area, EBA: extrastriate body area, PPA: parahippocampal place area, MPA: medial place area, OPA: occipital place area.

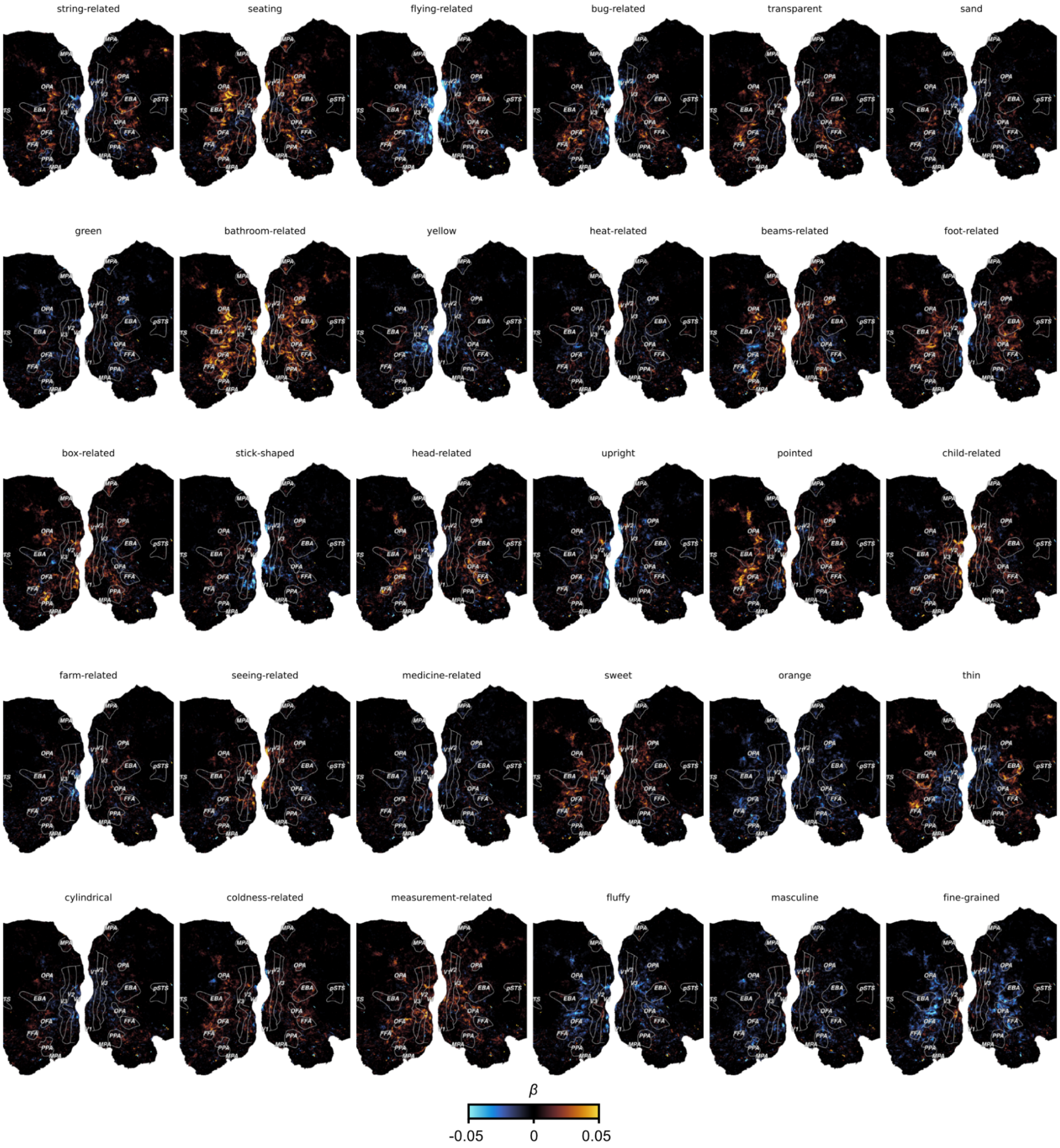


Extended Data Figure 2. **Dimension tuning maps 37-66 for Subject 1**. Colors indicate regression weights for each dimension predictor from the parametric modulation encoding model.

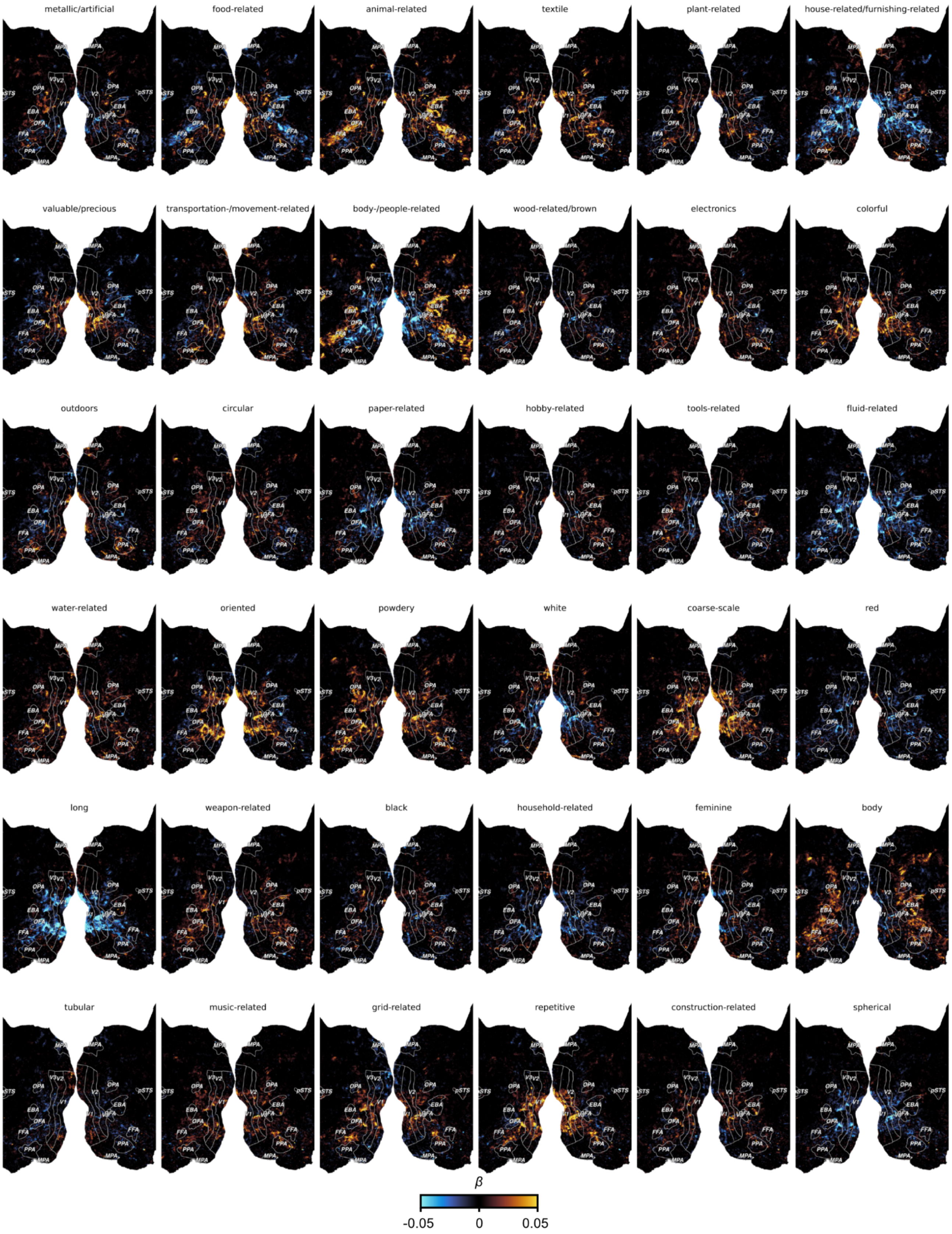


Extended Data Figure 3. **Dimension tuning maps 1-36 for Subject 2**. Colors indicate regression weights for each dimension predictor from the parametric modulation encoding model.

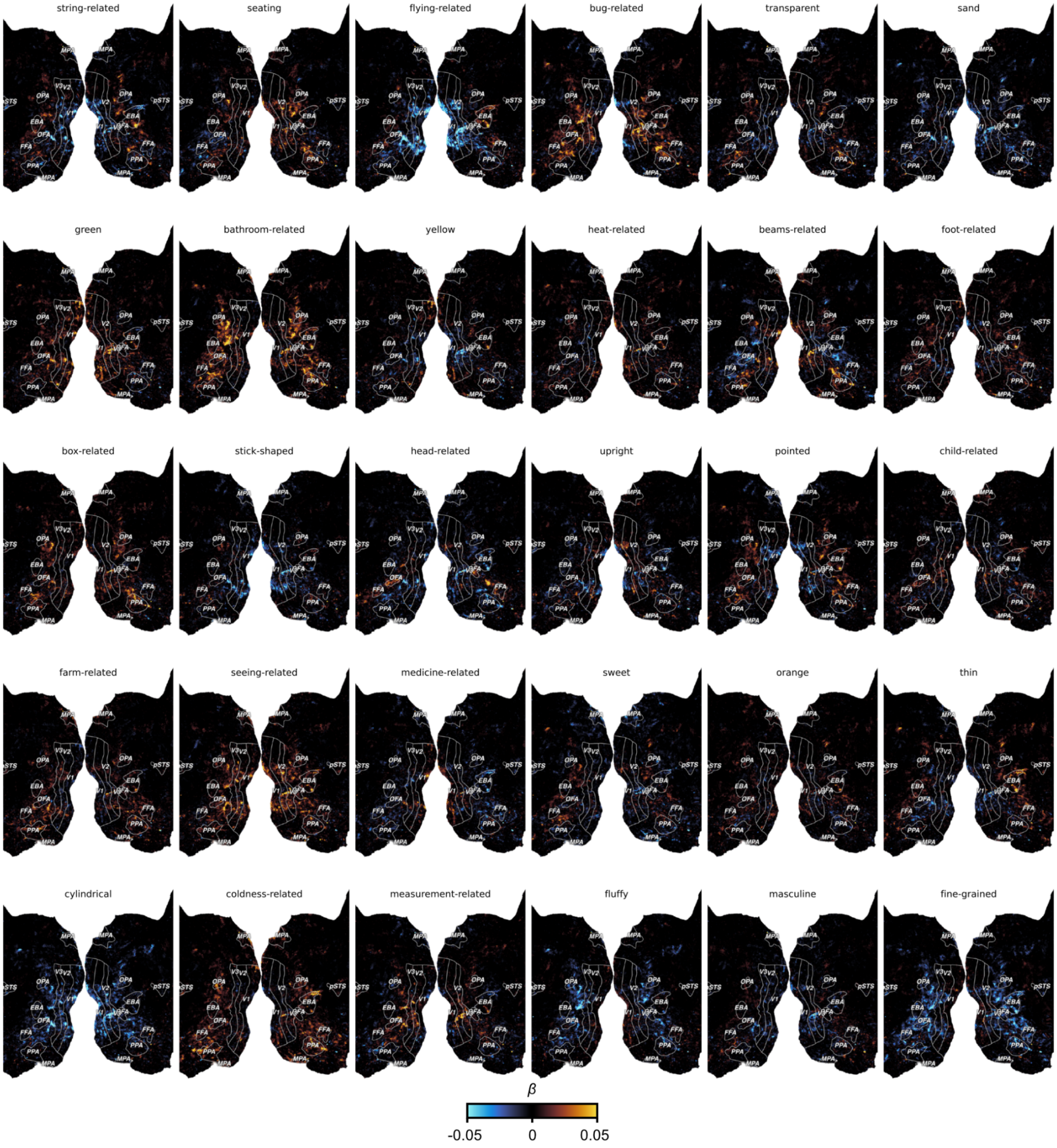


Extended Data Figure 4. **Dimension tuning maps 37-66 for Subject 2**. Colors indicate regression weights for each dimension predictor from the parametric modulation encoding model.

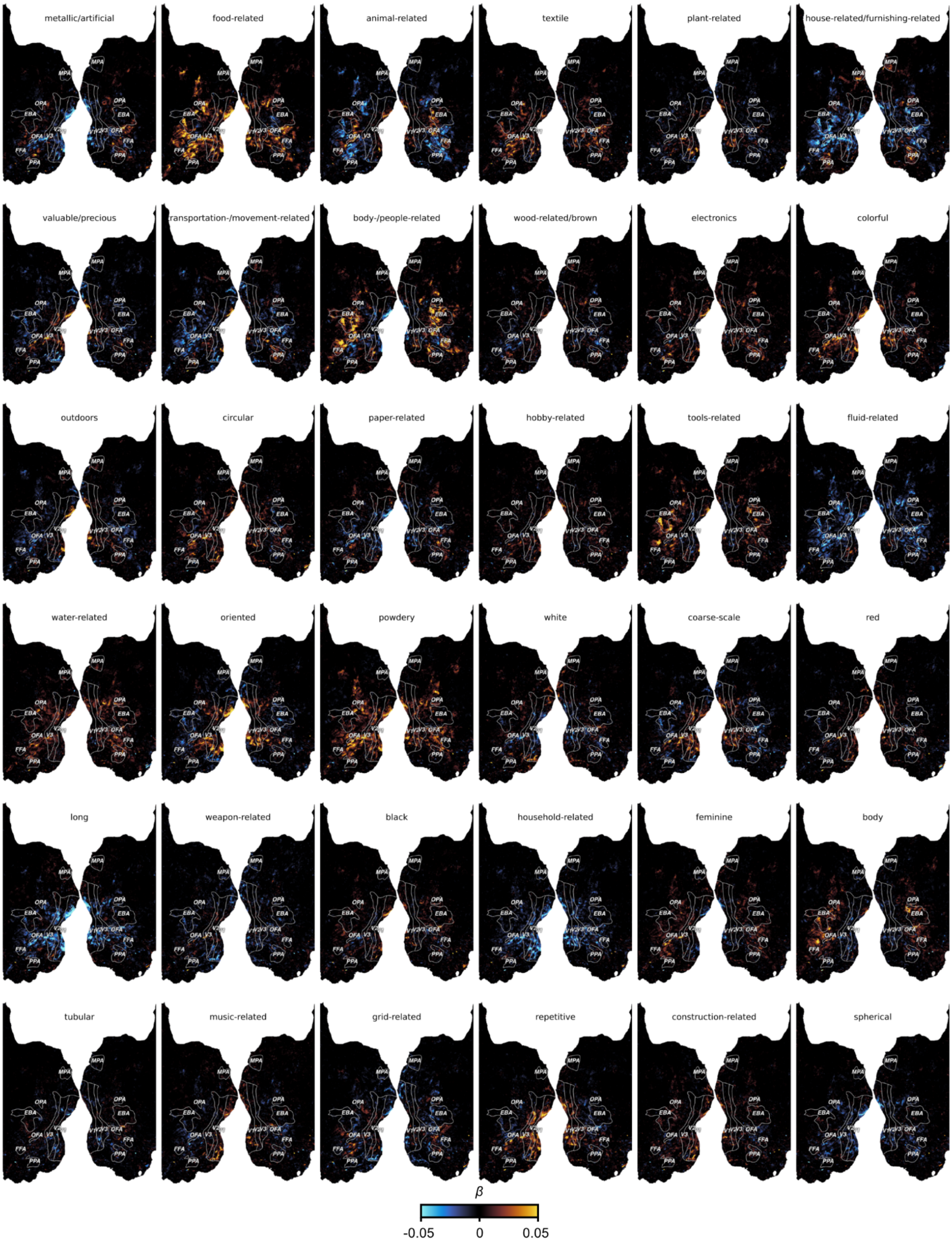


Extended Data Figure 5. **Dimension tuning maps 1-36 for Subject 3**. Colors indicate regression weights for each dimension predictor from the parametric modulation encoding model.

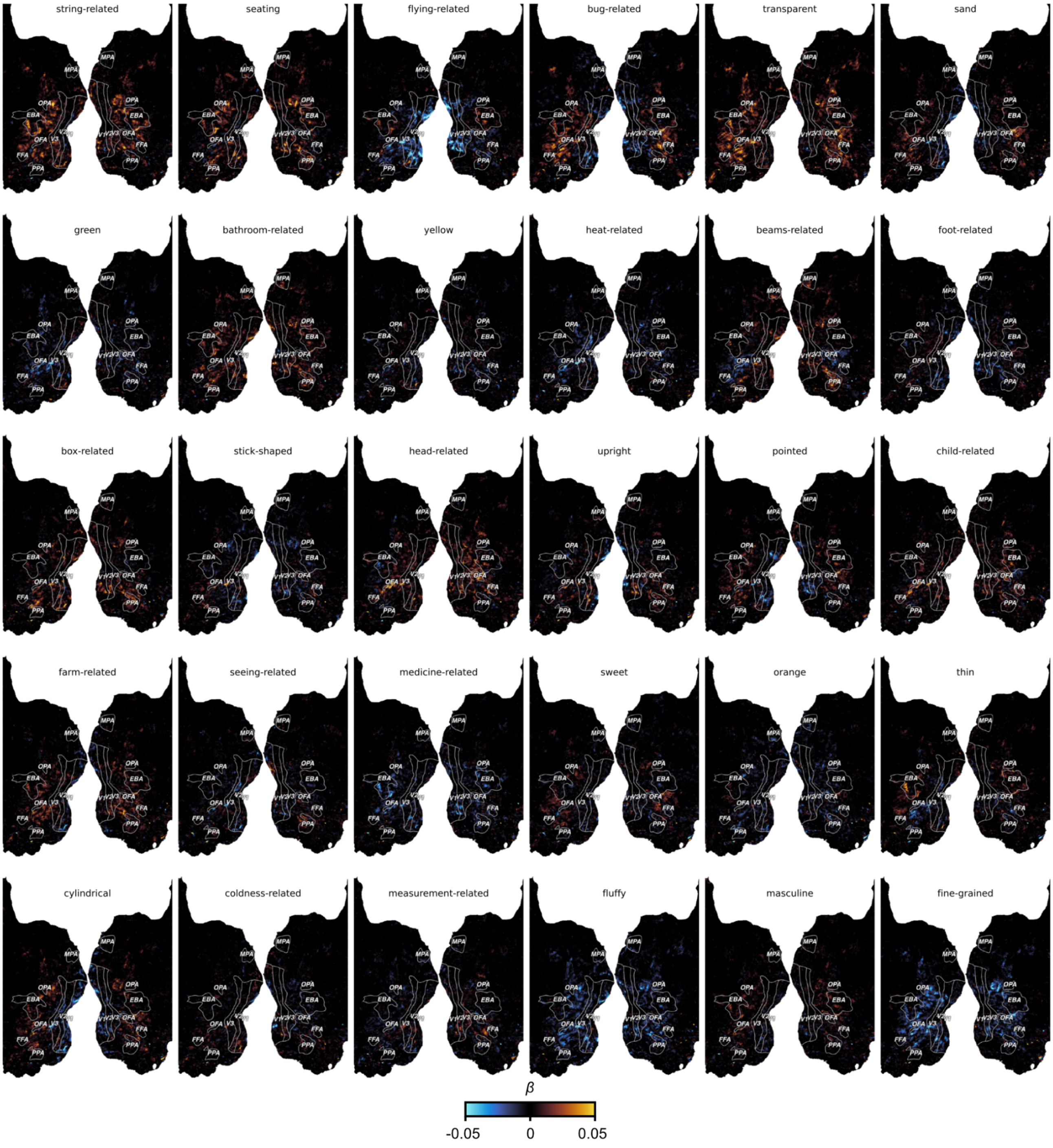


Extended Data Figure 6. **Dimension tuning maps 37-66 for Subject 3**. Colors indicate regression weights for each dimension predictor from the parametric modulation encoding model.
